# A conjugative plasmid exploits flagella rotation as a cue to facilitate its transfer

**DOI:** 10.1101/2024.07.18.604039

**Authors:** Saurabh Bhattacharya, Michal Bejerano-Sagie, Miriam Ravins, Liat Zeroni, Prabhjot Kaur, Venkadesaperumal Gopu, Ilan Rosenshine, Sigal Ben-Yehuda

## Abstract

Conjugation-mediated DNA delivery is the primary mode for antibiotic resistance spread; yet, molecular mechanisms regulating the process remain largely unexplored. While conjugative plasmids typically rely on solid surfaces to facilitate donor-to-recipient proximity, the pLS20 conjugative plasmid, prevalent among Gram-positive *Bacillus* spp., uniquely requires fluid environments to motivate its transfer. Here we unveiled that pLS20, carried by *B. subtilis*, induces adhesin-promoted multicellular clustering, which can accommodate various species, offering a stable platform for DNA delivery in liquid milieu. We further discovered that induction of pLS20 promoters, governing crucial conjugative genes, hinges on the presence of donor cell flagella, the major bacterial motility organelle. Moreover, pLS20 regulatory circuit is strategically integrated into a mechanosensing signal transduction pathway responsive to flagella rotation, harnessing propelled flagella to activate conjugation genes exclusively during the host motile phase. This flagella-conjugation coupling strategy, provides the plasmid with the benefit of disseminating into remote destinations, infiltrating new niches.

## Introduction

Conjugation, discovered by Lederberg and Tatum in 1946, is a key mechanism of horizontal gene transfer within natural bacterial communities, allowing a unidirectional transfer of genetic material from a donor to a recipient in a contact-dependent manner^1^. Conjugative plasmids, carrying genetic information essential for their autonomous transfer, play a central role in acquisition and dissemination of antibiotic resistance and virulence factors, thereby contributing to bacterial endurance and pathogenesis^e.g.2–4^. Successful conjugative transmission necessitates a tight association between donor and recipient bacteria, known as mating pair formation^e.g.5,6^. While Gram-negative conjugation systems typically achieve this intimate interaction through extracellular filaments, designated pili^e.g.7–10^, the mechanisms underlying mating pair formation in Gram-positives remain largely elusive, as their conjugation systems appear to lack a pilus or a similar structure^e.g.4^. Nonetheless, fundamental aspects of conjugation are conserved across Gram-positive and -negative bacteria. Both mechanisms are typified by converting conjugative plasmid DNA from double-stranded into a single-stranded form, a process driven by a plasmid- encoded relaxase. The resulting single-stranded DNA (ssDNA), protected by plasmid-encoded ssDNA binding protein (SSB), is directed to the transferosome, a membrane-associated mating complex housing a type IV secretion system (T4SS). Through this apparatus, the plasmid ssDNA is delivered to a recipient cell (transconjugant), providing a template for generating a double- stranded plasmid^e.g.4,10–13^.

The 65 Kb conjugative plasmid pLS20, initially isolated from the Gram-positive *Bacillus subtilis* (*B. subtilis*) *natto* strain^14^, possesses transmission capacity to other *Bacilli*, including the pathogenic species *B. anthracis* and *B. cereus*, highlighting its potential to serve as a vehicle for interspecies genetic exchange^15–17^. In fact, pLS20 serves as the archetype plasmid to a family of at least 35 members, dispersed across various *Bacillus* species residing in diverse ecological niches^18^. It carries a gene cluster encoding the T4SS machinery, SSB, relaxase, as well as additional essential conjugation functions^19–22^. The primary conjugation promoter P_C_ is repressed by Rco, maintaining the default "OFF" state of conjugation. In parallel, an anti-repressor, termed Rap, is held inactive through binding to a short quorum-sensing signaling peptide, Phr, encoded by the plasmid. The transition to an "ON" state is facilitated by a decrease in Phr concentration, and the consequent liberation of Rap, which counteracts Rco, thus promoting conjugation^23–26^.

Intriguingly, unlike most known conjugative systems that rely on a solid surface for efficient DNA delivery, pLS20 conjugation is typically executed in liquid medium, when the bacterium is highly motile^27^. Notably, *B. subtilis* exhibits two distinct lifestyles: a motile mode dominated by flagellated cells and a sessile mode marked by long cell chains, often devoid of flagella^28,29^. The switch between these modes is governed, among others, by the phosphodiesterase YmdB, with cells lacking *ymdB* displaying increased flagellation but are deficient in forming biofilms and intercellular membranous nanotube bridges^29–31^. Flagella biogenesis is initiated through the assembly of the flagellar type III secretion system, encoded by *fliOPQR flhBA* genes, collectively termed *CORE*^32–35^. This assembly, along with subsequent basal body and hook formation, triggers the secretion of FlgM, an anti-sigma factor inhibiting the activity of the flagellar sigma factor, SigD. Upon release of SigD from FlgM inhibition, the expression of various flagellar genes, including *hag*, encoding the major filament subunit, is induced, leading to the formation of the flagellar filament on the cell surface^34,36–39^.

Here we explore the enigmatic connection between *B. subtilis* motility and conjugation. We reveal that flagella are essential for effective conjugation in liquid media, by serving as a mechanosensory device prompting the transcription of genes essential for pLS20 conjugation in a subset of cells. Successively, these donor cells acquire the ability to organize mating clusters, fostering proximity between donor and recipient bacteria, establishing a robust foundation for mating pair formation, ultimately facilitating fruitful plasmid transfer.

## Results

### Visualization of conjugation revealed the formation of mating clusters

The dependency of pLS20 conjugation on *B. subtilis* motile lifestyle^27^, prompted us to explore how donor and recipient cells are coupled to fasten plasmid delivery in fluid surroundings. To track conjugation events at a single cell resolution, we initially employed pLS20, modified to include *gfp* driven by P_IPTG_ promoter, though GFP expression kinetics in the transconjugants was relatively slow (Figure S1A-S1B). To improve transconjugant detection, the native pLS20 *ssb* gene, encoding ssDNA binding protein^23^, one of the earliest proteins to be robustly expressed in transconjugants^40,41^, was fused to YFP. The SSB-YFP fusion was fully functional in supporting conjugation (Figure S1C), and was manifested within donor bacteria as faint foci (Figure 1A). When donor cells carrying pLS20-*ssb-yfp* were mixed with mCherry-labeled recipients, bright prominent SSB-YFP foci subsequently emerged in transconjugant bacteria, as early as 60 minutes post mixing (Figure 1A and Figure S1A-S1B). The appearance of these foci reported conjugation events wherein pLS20 was successfully delivered to recipient bacteria, leading to production of SSB-YFP.

**Figure 1:**
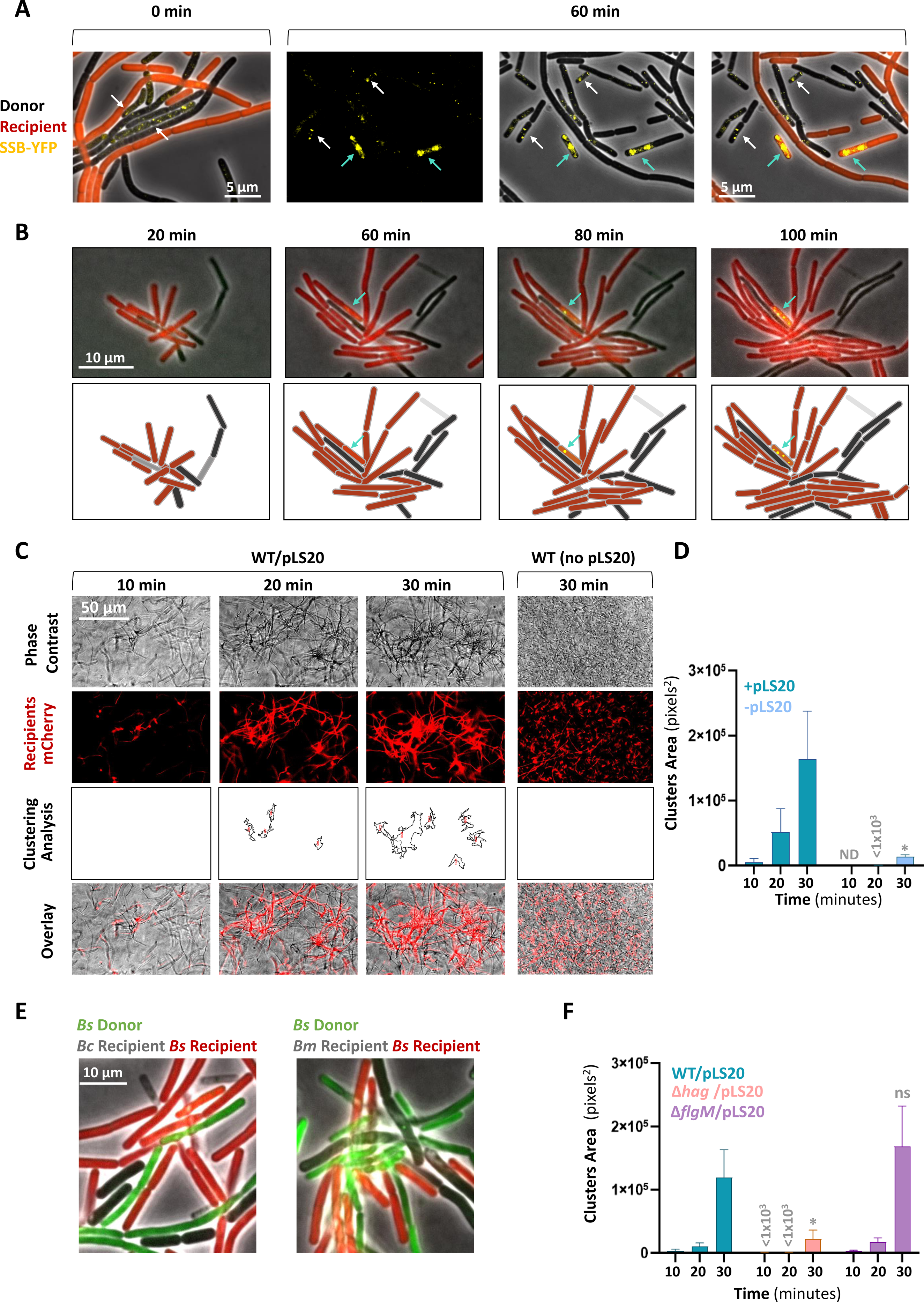
MC formation is promoted by pLS20 and donor flagella. **A.** Donor cells (dark) (SH347: WT/pLS20_cm_-*ssb-yfp*) were mixed with recipient cells (red) (BDR2637: *sacA::*P*_veg_-mCherry*) in 1:1 ratio (T=0 min), incubated for 60 minutes and visualized by fluorescence microscopy. Images were set to co-visualize faint (donor) and bright (transconjugant) SSB-YFP foci. Left panel: Shown is a representative overlay image of phase contrast (grey), with fluorescence from mCherry (red) and SSB-YFP (yellow) captured at 0 minutes. Right panels: Shown are representative images of fluorescence from (left to right): SSB-YFP (yellow), an overlay of phase contrast (grey) with fluorescence from SSB-YFP (yellow), and an overlay of phase contrast (grey), with fluorescence from mCherry (red) and SSB-YFP (yellow). White arrows highlight donor cells expressing faint SSB-YFP foci. Cyan arrows highlight transconjugants expressing bright SSB-YFP foci. A representative experiment out of 3 independent biological repeats. **B.** Donor cells (dark) (SH347: WT/pLS20_cm_-*ssb-yfp*) were mixed with recipient cells (red) (BDR2637: *sacA::*P*_veg_-mCherry*) in 1:1 ratio, placed over semi-solid (0.6%) LB agarose pad, and visualized by time lapse fluorescence microscopy. Upper panels: overlay images of phase contrast (grey) with fluorescence from mCherry (red) and SSB-YFP (yellow), captured at the indicated time points. Lower panels: Schematics depicting cell layout of the corresponding upper panels with a transconjugant cell highlighted by an arrow. **C.** Donor cells (dark) (SH337: WT/pLS20_cm_) were mixed with recipient cells (red) (BDR2637: *sacA::*P*_veg_-mCherry*) in 1:1 ratio, and the formation of MCs in liquid medium was followed using digital wide-field microscopy. A strain lacking pLS20_cm_ (PY79: WT) was used as a control. Shown are representative images captured at the indicated time points after mixing: phase contrast (grey), fluorescence from mCherry-labeled recipients (red), computed clustering analysis (outlines), and overlay of phase contrast and mCherry fluorescence. A representative experiment out of 3 independent biological repeats. **D.** Quantification of the clustering analysis shown in (C). The presence or absence of pLS20_cm_ is indicated. For each mixture, triplicates were analyzed in parallel, and total area of clusters larger than 500 pixels^2^ was calculated. Data is presented as average values and SEM. Statistical significance was calculated using two-way ANOVA. For t=30 min, P-values: (*) ≤ 0.05. ND-not detected. **E.** *B. subtilis* (*Bs*) donor cells (green) (SH501: *sacA*::P_33_-*gfp*/pLS20_cm_) were mixed with *B. subtilis* recipient cells (red) (BDR2637: *sacA::*P*_veg_-mCherry*) and *B. cereus* (*Bc*) (dark) (left panel) or *B. megaterium* (*Bm*) (dark) (right panel) cells in 1:1:1 ratio, placed over semi-solid (0.6%) LB agarose pad for 45 minutes and visualized by fluorescence microscopy. Shown are overlay images of phase contrast (grey), with fluorescence from mCherry (red) and GFP (green). **F.** Donor strains (dark): WT (SH337), Δ*hag* (SH443), or Δ*flgM* (SH496) harboring pLS20_cm_, were mixed with recipient cells (red) (BDR2637: *sacA::*P*_veg_-mCherry*) in 1:1 ratio, and the formation of MCs in liquid medium was followed using digital wide-field microscopy. Shown is quantification of the clustering analysis at the indicated time points after mixing. Representative images for time point 30 minutes are displayed in Figure S2A. For each mixture, triplicates were analyzed in parallel, and total area of clusters larger than 500 pixels^2^ was calculated. Data is presented as average values and SEM. Statistical significance between WT and each mutant was calculated using two-way ANOVA. For t=30 min, P-values: (ns) not significant ≥ 0.05, (*) ≤ 0.05. A representative experiment out of 3 independent biological repeats.

Consequently, we developed a microscopy-based conjugation assay under semi-liquid conditions, allowing to follow the kinetics of conjugation events by time lapse microscopy. Intriguingly, promptly after mixing donor and recipient bacteria in a 1:1 ratio, cell clusters emerged, comprising a few donors surrounded by numerous recipients. These cell assemblages remained stable and expanded over time both through cell division and the incorporation of additional bacteria (Figure 1B). Approximately 60 to 80 minutes post co-incubation, recipient transconjugants expressing SSB-YFP appeared adjacent to donor bacteria; hence, reporting effective conjugation events (Figure 1B). Notably, transconjugants were predominantly cluster- associated (∼85%) (Figure S1D), suggesting that donor cells induce the formation of multi-cellular "mating-clusters" (MCs), providing a platform for enhancing conjugation. To further examine this notion, we designed an assay for quantitative monitoring of MC formation kinetics after mixing donors with mCherry-labeled recipient cells, using digital wide-field microscopy. MCs, dominated by recipient bacteria, were typically evident 20 minutes post co-incubation, becoming more pronounced over time (Figure 1C-1D). Importantly, MC development was pLS20-dependent, strongly linking the phenomenon to the conjugation process (Figure 1C-1D). Since pLS20-like plasmids are prevalent in *Bacilli*^18^, we investigated whether species other than *B. subtilis* could be recruited to pLS20-derived MCs. Therefore, MC formation was visualized within mixed populations comprising differentially-labeled *B. subtilis* donor and recipient cells, alongside unlabeled *B. cereus* or *B. megaterium* cells. Remarkably, cells from both species were observed associating with *B. subtilis* MCs, and were often adjacent to potential *B. subtilis* donors (Figure 1E).

Taken together, during conjugation, pLS20-induced MCs are assembled, containing a few donors outnumbered by potential recipients, which could belong to the same or different species. These MCs appear to provide a relatively static hub for promoting intra- and inter- species mating pair formation in liquid conditions.

### Donor-produced flagella are required for conjugation

To address whether bacterial motility impacts MC establishment, a mutant lacking the flagellum filament (Δ*hag*) and a hyper-flagellated mutant, overexpressing the flagellar genes (Δ*flgM*)^34,39^, both carrying pLS20, were tested for their ability to assemble MCs. Surprisingly, the kinetics of MC formation was largely impeded in the non-motile Δ*hag* donor, whereas the hyper-motile Δ*flgM* showed enhanced clustering capacity (Figure 1F and Figure S2A). Nonetheless, the lack of flagella in the recipient did not perturb MC formation (Figure S2B), excluding that flagella entangling or adhesion properties^42–44^ are involved in this process. We then explored whether the dependency of MC formation on flagella correlates with respective conjugation efficiencies, utilizing the SSB-YFP reporter. No discernible conjugation events were monitored for Δ*hag* donor, whereas the Δ*flgM* donor yielded elevated transconjugant levels compared to WT (Figure 2A-2B). These results indicate that MC formation and the subsequent pLS20 conjugation are promoted by donor flagella.

**Figure 2:**
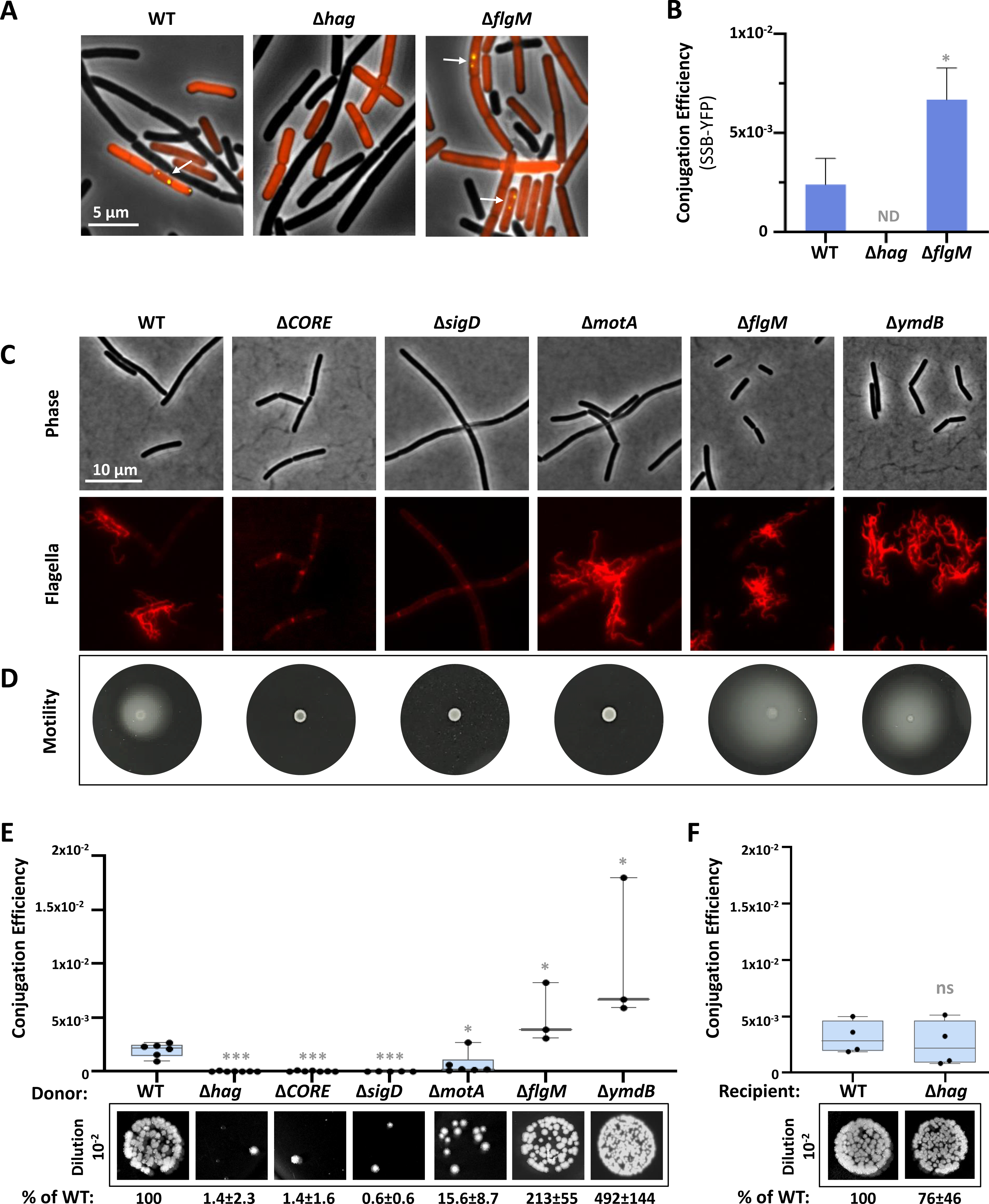
Functional flagella facilitate conjugation. **A.** Donor strains (dark): WT (SH347), Δ*hag* (MBS20), or Δ*flgM* (SH561), harboring pLS20_cm_-*ssb-yfp*, were mixed with recipient cells (red) (BDR2637: *sacA::*P*_veg_-mCherry*) in 1:1 ratio, incubated for 1 hour and visualized by fluorescence microscopy. Shown are overlay images of phase contrast (grey), with fluorescence from mCherry (red) and SSB-YFP (yellow) of the indicated strains. Arrows highlight transconjugant cells. **B.** Conjugation efficiencies of the donors described in (A) were calculated as the number of transconjugants [mCherry (red) cells displaying SSB-YFP (yellow) foci]/number of recipients [mCherry (red) cells]. Data is presented as average values and SEM of at least seven fields from a representative experiment out of 3 independent biological repeats. Statistical significance between WT and each mutant was calculated using one-way ANOVA. P-values: (*) ≤ 0.05. ND-not detected. **C.** The following strains: WT (DS1895), Δ*CORE* (SH411), Δ*sigD* (SH415), Δ*motA* (SH409), Δ*flgM* (SH408), and Δ*ymdB* (SH101), harboring modified flagellin *hag*^T209C^ (*amyE*::P*_hag_-hag*^T209C^), were grown in liquid LB, flagellin was stained with Alexa Fluor 594 C_5_ maleimide, and cells were visualized by fluorescence microscopy. Shown are images of phase contrast (grey) (upper panels) and corresponding stained flagella (red) (lower panels). **D.** The following strains: WT (PY79), Δ*CORE* (SH9), Δ*sigD* (IB97), Δ*motA* (SH419), Δ*flgM* (SH495) and Δ*ymdB* (GB61) were grown to mid-logarithmic phase and subjected to motility assay by spotting onto LB plates supplemented with 0.3% agar. Shown are images captured after 7 hours of incubation. **E.** Donor strains: WT (SH337), Δ*hag* (SH443), Δ*CORE* (SH352), Δ*sigD* (PK143), Δ*motA* (SH423), Δ*flgM* (SH496), or Δ*ymdB* (SH442) harboring pLS20_cm_ were mixed with recipient cells (SH345: *sacA::kan*) in 1:1 ratio, incubated for 20 minutes, and serial dilutions were spotted either on LB agar containing chloramphenicol and kanamycin or solely kanamycin, selecting for transconjugants and recipients, respectively. Upper panel: Conjugation efficiencies calculated as number of transconjugants/number of recipients. For each strain, at least 3 independent biological repeats were conducted. Data are shown as box plot graphs. The box is determined by the 25^th^ and 75^th^ percentiles, and whiskers are determined by min and max; the line in the box indicates the median. Statistical significance between WT and each mutant was calculated using unpaired t-tests. P-values: (*) ≤ 0.05, (***) ≤ 0.001. Lower panels: Representative images of spotted conjugation mixtures (10^-2^ dilution) over LB agar containing chloramphenicol and kanamycin, selecting for transconjugants. Indicated conjugation efficiencies were calculated as % of WT conjugation efficiency. Presented are average values and SD of at least 3 independent biological repeats. **F.** Donor cells (SH337: WT/pSL20_cm_) were mixed with WT (SH345: *sacA::kan*) or Δ*hag* (MBS11: *Δhag, amyE*::P_IPTG_-*gfp-kan*) recipient cells in 1:1 ratio, incubated for 20 minutes, and serial dilutions were spotted either on LB agar containing chloramphenicol and kanamycin or solely kanamycin, selecting for transconjugants and recipients, respectively. Upper panel: Conjugation efficiencies calculated as number of transconjugants/number of recipients. For each strain, at least 3 independent biological repeats were conducted. Data are shown as box plot graphs. The box is determined by the 25^th^ and 75^th^ percentiles, and whiskers are determined by min and max; the line in the box indicates the median. Statistical significance was calculated using paired t-tests. P-values: (ns) not significant ≥ 0.05. Lower panels: Representative images of spotted conjugation mixtures (10^-2^ dilution) over LB agar containing chloramphenicol and kanamycin, selecting for transconjugants. Indicated conjugation efficiencies were calculated as % of WT conjugation efficiency. Presented are average values and SD of at least 3 independent biological repeats.

To substantiate the link between the presence of flagella and conjugation, we quantified the rate of emergence of transconjugant colony-forming units (CFUs), in various donor flagellar mutants, concomitantly with flagella visualization and motility (Figure 2C-2E). In agreement with the microscopy-based assay, the non-motile Δ*hag* cells were severely conjugation deficient (Figure 2E). We further tested additional non-motile mutants perturbed in flagella biogenesis: Δ*CORE* deleted for the genes encoding the flagellar export apparatus, and Δ*sigD*, lacking the flagellar sigma factor^34,36^ (Figure 2C-2D). Both mutants were strongly attenuated in pLS20 conjugation (Figure 2E). Moreover, in line with the microscopy findings, the hyper-motile mutant, Δ*flgM*, exhibited increased conjugation levels, that were further boosted in Δ*ymdB* mutant overexpressing flagella^29^ (Figure 2C-2E). Interestingly, a non-motile mutant lacking *motA*, which assembles paralyzed flagella^45^, showed decreased conjugation ability (Figure 2C-2E), suggesting that rotating flagella is required for the process.

To inspect whether the necessity of flagella for conjugation is solely a donor property or extends to the recipient, conjugation was conducted by mating a WT donor with either WT or Δ*hag* recipients. Similar numbers of transconjugants were obtained for both recipients, signifying that the need for flagella is a donor-specific demand (Figure 2F), consistent with the flagella dispensability for recipient bacteria during MC formation (Figure S2B). Collectively, our results indicate that functional rotating donor flagella collaborate with pLS20 factors to induce MC formation and conjugation occurrence.

### Flagella are essential for conjugation gene expression

To study the nature of the link between flagellum functionality and conjugation events, we assessed whether the necessity for flagella within donor cells could be bypassed by enhancing the transcription of conjugative genes. To achieve this, the anti-Rco conjugation repressor, RapA_pLS20_ (herein RapA)^23,26^, was ectopically overexpressed in flagellar mutants and assayed for conjugation. Indeed, ectopic expression of RapA effectively bypassed the conjugation defect in all tested flagellar mutants (Figure 3A), hinting that flagella might play a role in augmenting the transcription of conjugation genes.

**Figure 3:**
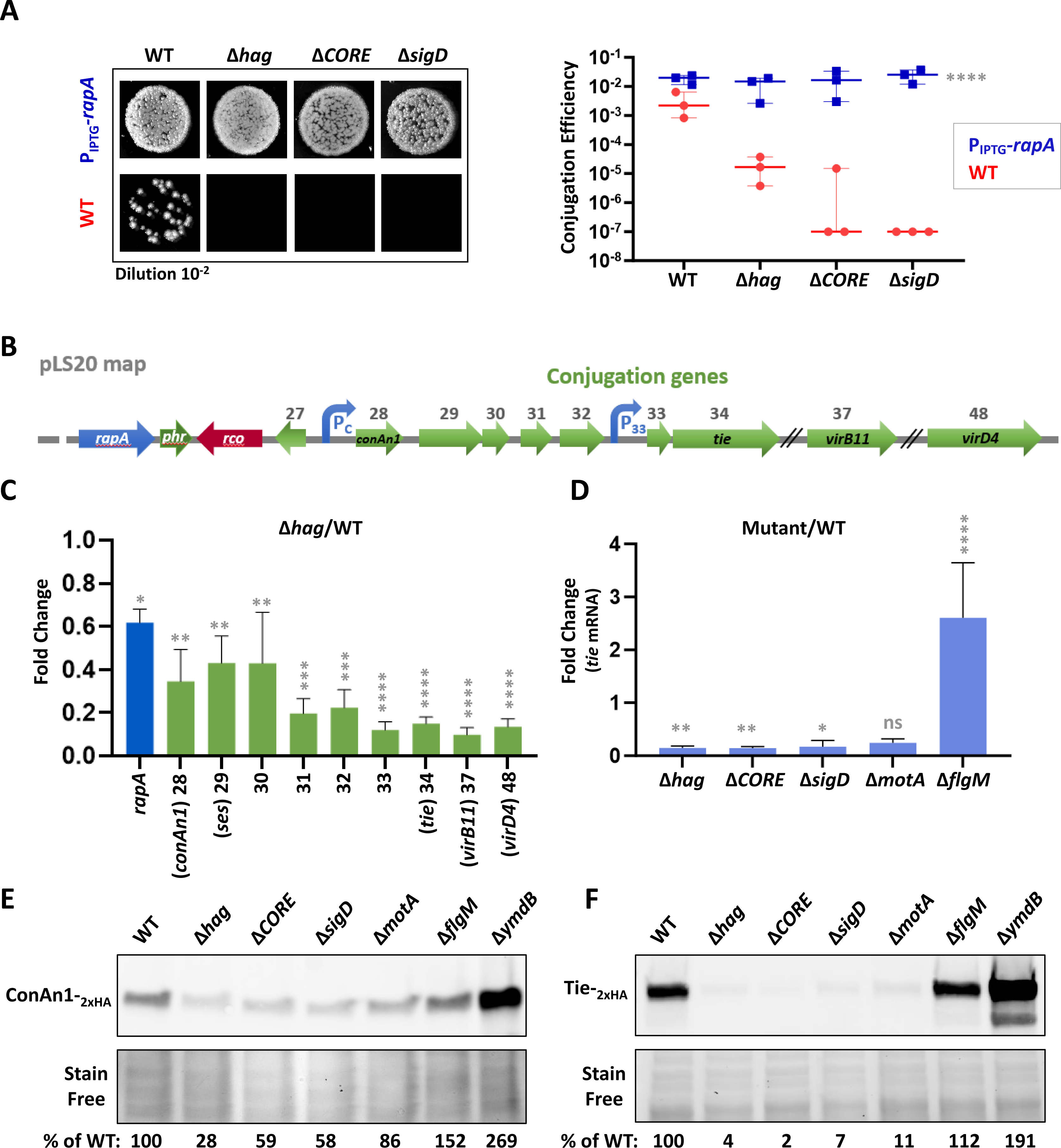
Flagella affect conjugation gene expression. **A.** Donor strains: WT (SH337), Δ*hag* (SH443), Δ*CORE* (SH352), and Δ*sigD* (PK143) harboring pLS20_cm_, and donor strains: WT (SH450), Δ*hag* (SH453), Δ*CORE* (SH451), and Δ*sigD* (SH452), harboring both pLS20_cm_ *and amyE*::P_IPTG_-*rapA*_pLS20_, were grown in the presence of IPTG (0.1 mM) and mixed with recipient cells (SH345: *sacA::kan*) in 1:1 ratio. Conjugation mixes were incubated for 20 minutes and serial dilutions were spotted either on LB agar containing chloramphenicol and kanamycin or solely kanamycin, selecting for transconjugants and recipients, respectively. Left panel: Representative images of spotted conjugation mixtures (10^-2^ dilution) over LB agar containing chloramphenicol and kanamycin, selecting for transconjugants. Right panel: Conjugation efficiencies calculated as number of transconjugants/number of recipients, in the presence (blue squares) or absence (red circles) of ectopically expressed *rapA*. Data is presented as scatter dot plot with a line indicating the median with range, of 3 independent biological repeats. Statistical significance was calculated using two-way ANOVA. P-values of P_IPTG_-*rapA*_pLS20_ vs WT: (****) ≤ 0.0001. **B.** Schematic depicting the conjugation operon of pLS20. Regulatory elements controlling pLS20 operon expression, as well as known encoded genes, are highlighted. **C.** RNA was isolated from WT (SH337) and Δ*hag* (SH443) donor cells harboring pLS20_cm_ and mRNA levels of the indicated pLS20 genes were determined using qRT-PCR. Shown are relative transcript levels of the indicated genes in Δ*hag* compared to the WT. Data is presented as average values and SEM of at least 3 independent biological repeats. Statistical significance was calculated using one-way ANOVA. P-values: (*) ≤ 0.05, (**) ≤ 0.005, (***) ≤ 0.0005, (****) ≤ 0.0001. **D.** RNA was isolated from WT (SH337), Δ*hag* (SH443), Δ*CORE* (SH352), Δ*sigD* (PK143), Δ*motA* (SH423) and Δ*flgM* (SH496) donor strains harboring pLS20_cm_, and mRNA level of *tie* was determined using qRT-PCR. Shown are relative transcript levels of *tie* in the indicated mutant strains compared to the WT. Data is presented as average values and SEM of at least 3 independent biological repeats. Statistical significance was calculated using one-way ANOVA. P-values: (ns) not significant ≥ 0.05, (*) ≤ 0.05, (**) ≤ 0.005, (****) ≤ 0.0001. **E.** Whole cell lysates were extracted from donor strains: WT (MR13), Δ*hag* (LZ4), Δ*CORE* (LZ5), Δ*sigD* (LZ6), Δ*motA* (LZ7), Δ*flgM* (LZ9) and Δ*ymdB* (LZ8), harboring pLS20_cm_-*conAn1_-_*_2xHA_, and subjected to western blot analysis using anti-HA antibodies (upper panel). Stain-free total protein analysis is presented for comparison (lower panel). ConAn1_-2xHA_ expression levels, normalized to stain-free total protein were calculated as % of WT. Shown is a representative experiment out of 3 independent biological repeats. **F.** Whole cell lysates were extracted from donor strains: WT (SH461), Δ*hag* (MBS24), Δ*CORE* (MBS26), Δ*sigD* (MBS25), Δ*motA* (SH481), Δ*flgM* (LZ16) and Δ*ymdB* (MBS23), harboring pLS20_cm_-*tie_-_*_2xHA_ and subjected to western blot analysis using anti-HA antibodies (upper panel). Stain-free total protein analysis is presented for comparison (lower panel). Tie_-2xHA_ expression levels, normalized to stain-free total protein were calculated as % of WT. Shown is a representative experiment out of 3 independent biological repeats.

The potential flagella-conjugation gene expression axis, was further examined by conducting quantitative Reverse Transcriptase PCR (qRT-PCR) on select pLS20 genes, comparing their transcription between WT and Δ*hag* mutant. A modest, yet significant, reduction in the RNA levels of *rapA*, along with ORFs 28 to 30, was detected in the Δ*hag* mutant (Figure 3B-3C). Furthermore, a sharper drop in the RNA amounts of ORF33 and onwards was evident, including transcripts of key conjugation genes such as *tie* (ORF34), encoding a putative adhesin, as well as *virB11* and *virD4*, encoding T4SS components^23,46^ (Figure 3B-3C). This gene expression dependency on flagella was emphasized by assessing *tie* transcription in various flagellar mutants, corroborating this phenomenon (Figure 3D).

To strengthen these findings, we surveyed the protein level of crucial conjugation factors within flagellar mutants. pLS20-encoded *conAn1* (ORF28), an anti-termination factor^47^, and *tie* were HA-tagged, and their production examined. ConAn1-_2xHA_ levels were markedly reduced in the flagella-deficient mutants, while exhibiting increased synthesis in the hyper-flagellated strains (Figure 3E). Consistent with the RNA measurements, flagella exerted an even stronger effect on Tie-_2xHA_ expression (Figure 3D and 3F). Taken together, these results show that intact functional flagella are vital for expression of essential conjugation genes.

### Conjugation genes are activated in a subpopulation in a flagella-dependent manner

To directly test the flagella impact on conjugation gene expression, we monitored conjugation promoter activity. At first, the key conjugation promoter, P_C_^24,48^ (Figure 3B), was fused to *gfp*, and the reporter was inserted into the host chromosome. Further, given the relatively lengthy intergenic region between ORFs 32 and 33 and the significant decrease observed in RNA levels of downstream genes in the Δ*hag* mutant (Figure 3B-3C), the presence of an additional promoter in this location was investigated. We thus similarly fused this putative promoter to *gfp* and introduced the reporter into the host genome. The activity of the promoters was then evaluated in the absence or presence of pLS20. Fluorescence measurements revealed the existence of an active promoter, although weaker than P_C_, in the ORFs 32-33 intergenic region, termed P_33_ (Figure 4A-4B). While P_C_ activity was sharply inhibited in the presence of pLS20, that of P_33_ remained constant (Figure 4A-4B). Surprisingly, the absence or overexpression of flagella appeared to have only a marginal impact, if any, on the overall population fluorescence (Figure 4A-4B).

**Figure 4:**
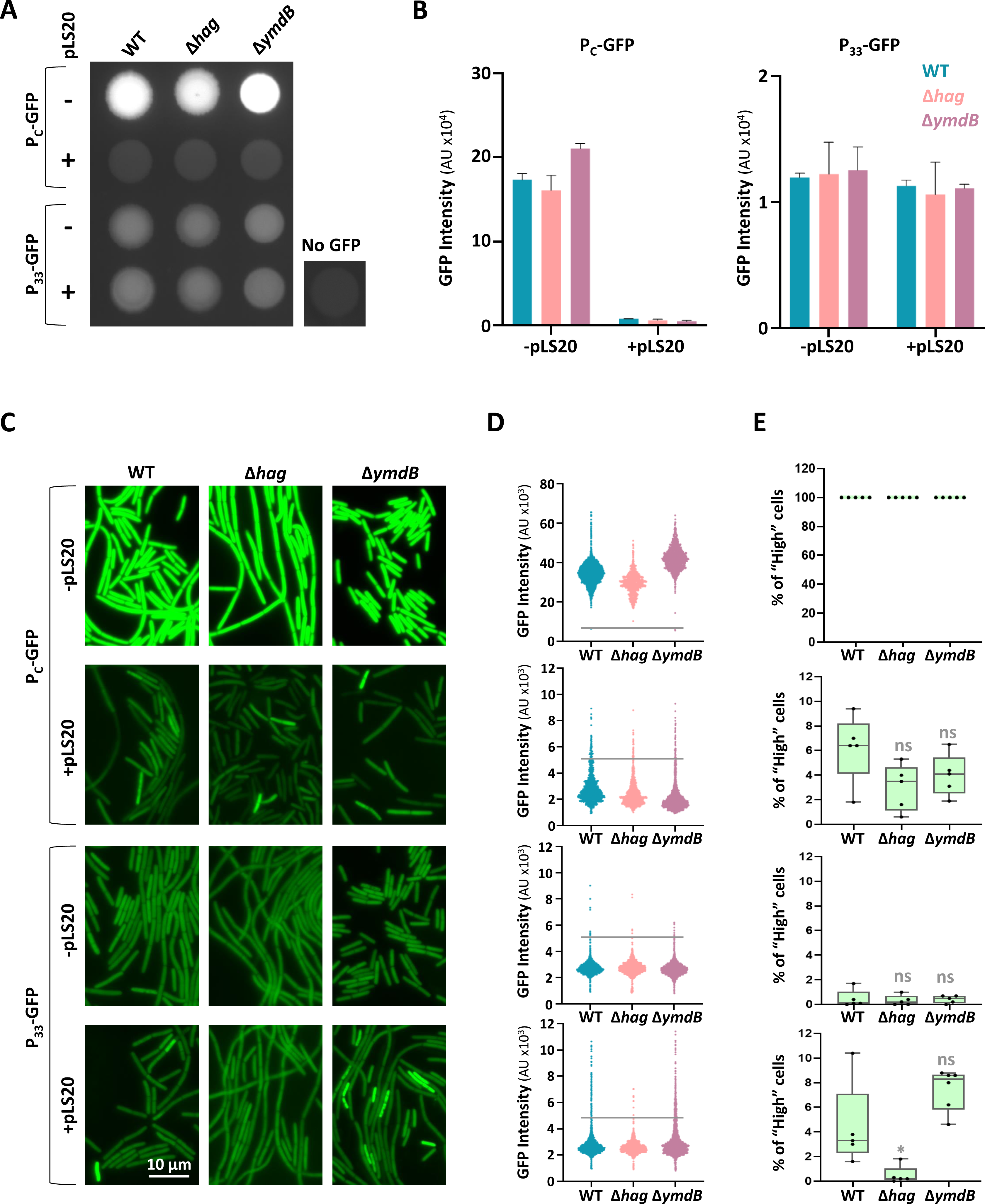
Conjugation promoters exhibit ON/OFF state. **A.** Strains harboring P_c_-*gfp* lacking pLS20_cm_, WT (SH437), Δ*hag* (SH458) and Δ*ymdB* (SH457), or carrying pLS20_cm_, WT (SH444), Δ*hag* (SH448) and Δ*ymdB* (SH447), and strains harboring P_33_-*gfp* lacking pLS20_cm_, WT (SH494), Δ*hag* (SH505) and Δ*ymdB* (SH504), or carrying pLS20, WT (SH501), Δ*hag* (SH510) and Δ*ymdB* (SH509), were grown in LB to OD_600_ 0.8 and spotted on an LB agar plate. Plates were incubated for 18 hours and visualized by fluorescence imaging system. A WT donor strain (SH337: WT/pLS20_cm_), lacking *gfp* (No GFP), served as a control. Shown is a representative experiment out of 3 independent biological repeats. **B.** Strains in (A) were grown in LB to OD_600_ 0.8, and GFP fluorescence was measured by a fluorescence microplate reader. Shown are normalized GFP intensities in arbitrary units (AU) from P_c_-GFP (left panel), and from P_33_-GFP (right panel). Each bar represents an average value and SD of 3 independent biological repeats. **C.** Strains in (A) were grown in LB to OD_600_ 0.8, and visualized by fluorescence microscopy. Shown are representative images of GFP fluorescence (green) from P_c_-GFP (upper panels) and P_33_-GFP (lower panels). **D.** Population distribution of cells based on their GFP expression in (C) in arbitrary units (AU). Each dot represents one cell. n>2000 cells for each strain. Horizontal grey line indicates GFP intensity threshold (5000 AU). Shown is a representative experiment out of 3 independent biological repeats. **E.** Quantification of the percentage of bacterial cells with GFP fluorescence higher than 5000 AU, designated as "High" GFP expressing cells, in the indicated strains following fluorescence microscopy analysis displayed in (D). For each strain, at least 3 fields were quantified. Data are shown as box plot graphs. The box is determined by the 25^th^ and 75^th^ percentiles, and whiskers are determined by min and max; the line in the box indicates the median. Statistical significance between WT and each mutant was calculated using one-way ANOVA. P-values: (ns) not significant ≥ 0.05, (*) ≤ 0.05.

Considering that conjugation is a relatively rare event (∼2.5 x 10^-3^) (Figure 2B and 2E), we tracked the activity of the conjugation promoters at a single-cell resolution using fluorescence microscopy. In the absence of pLS20, both P_C_ and P_33_ showed uniform levels of GFP expression, irrespective of flagella presence, with P_C_ exhibiting a significantly higher activity (Figure 4C-4D). In the presence of pLS20, P_C_ expression was repressed in most bacteria; however, a small subpopulation emerged displaying elevated GFP expression (GFP "High" cells), which was slightly affected by flagella existence (Figure 4C-4E). Moreover, P_33_ exhibited a sharper flagella-dependent bimodal expression profile, with the "High" population hardly detected in the Δ*hag* mutant, yet manifested in the Δ*ymdB* hyper-flagellated mutant (Figure 4C-4E).

The strict dependency of P_33_ "High" subpopulation on flagella existence, implies that these cells express the conjugation machinery and could be predictive of forthcoming conjugation events. To investigate this prospect, we simultaneously visualized, P_33_-GFP activity in donor cells, conjugation events through the SSB-YFP reporter, and MC formation. Indeed, transconjugants, exhibiting prominent SSB-YFP foci, were predominantly located within MCs containing donor cells exhibiting "High" P_33_-GFP (Figure 5A), emphasizing that this small subpopulation drives conjugation.

**Figure 5:**
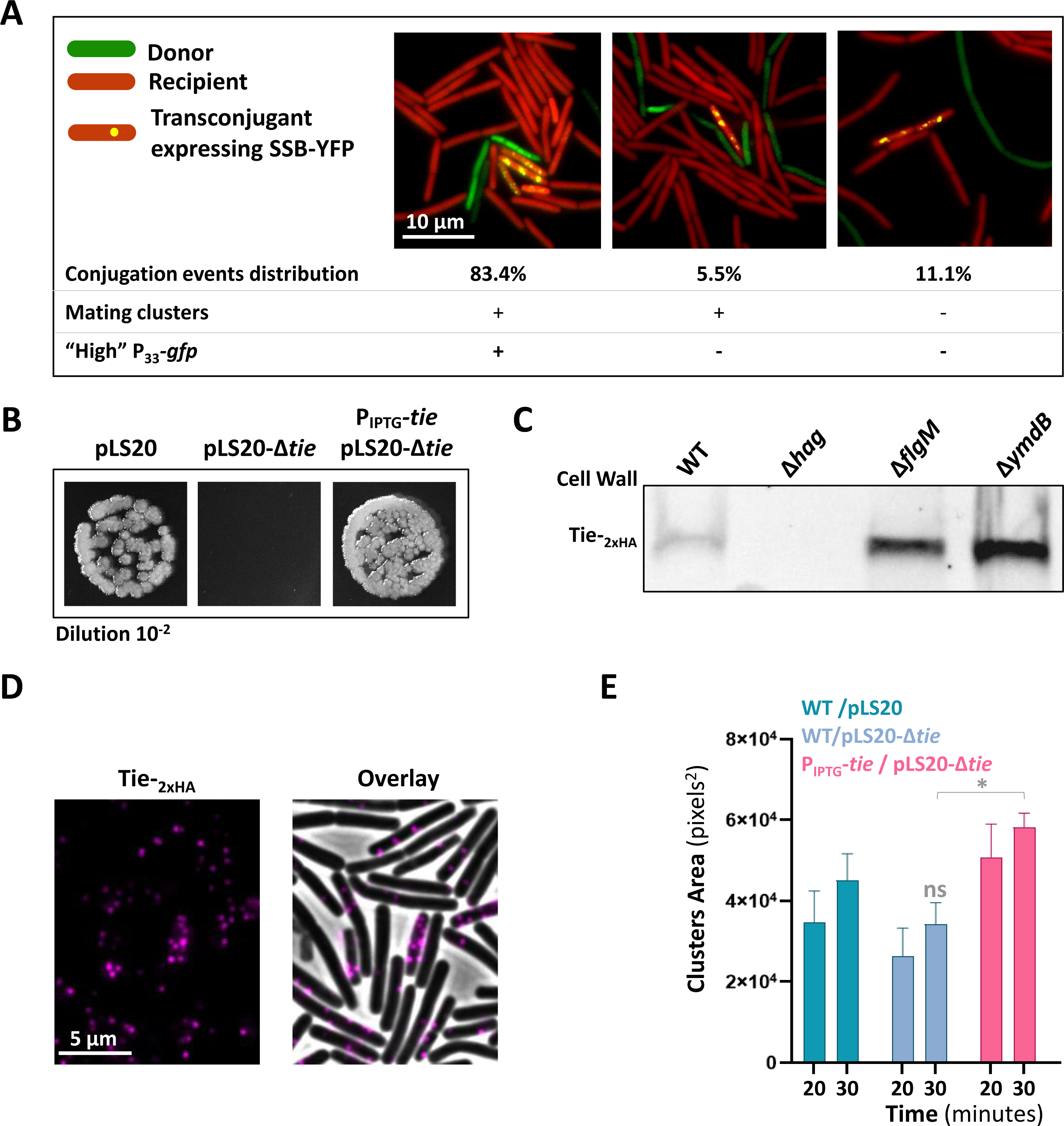
P_33_ "high" state coincides with MC formation and conjugation. **A.** Donor cells (green) harboring P*_33_*-*gfp* and pLS20_cm_-*ssb-yfp* (SH579) were mixed with recipient cells (red) (BDR2637: *sacA::*P*_veg_-mCherry*) in 1:1 ratio, placed over semi-solid (0.6%) LB agarose pads, incubated for 80 minutes, and visualized by fluorescence microscopy. Shown are overlay images of fluorescence from GFP (green), mCherry (red) and SSB-YFP (yellow). Images demonstrating transconjugant cells expressing SSB-YFP were categorized based on MC formation (should we quantify using an area threshold for phase?), and the presence of donor cells expressing "High" P*_33_*-*gfp* (>5000 arbitrary units). The distribution of conjugation events in each category is indicated in the table. Shown is a representative experiment out of 2 independent biological repeats. **B.** Donor strains: WT (SH337: WT/pLS20_cm_), Δ*tie* (SH483: WT/pLS20_cm_-Δ*tie*), and a *tie* complementing strain (SH485: *amyE*::P_IPTG_-*tie*/pLS20_cm_-Δ*tie*), were grown in the presence of IPTG (1 mM) and mixed with recipient cells (SH345: *sacA::kan*) in 1:1 ratio, and incubated for 20 minutes. Shown are images of spotted conjugation mixtures (10^-2^ dilution) over LB agar containing chloramphenicol and kanamycin, selecting for transconjugants. Representative images out of 3 independent biological repeats. **C.** Donor strains: WT (SH461), Δ*hag* (MBS24), Δ*flgM* (LZ16), and Δ*ymdB* (MBS23), harboring pLS20_cm_-*tie*_2xHA_ were grown to OD_600_ 0.8, cell-wall proteins were extracted using LiCl, and samples were subjected to western blot analysis using anti-HA antibodies. Shown is a representative experiment out of 3 independent biological repeats. **D.** Donor cells (SH461: WT/pLS20_cm_-*tie_-_*_2xHA_) were grown to OD_600_ 0.8 and visualized by immunofluorescence microscopy using primary anti-HA antibodies and Alexa647 conjugated secondary antibodies. Cells were not permeabilized before antibody treatment to enable the selective visualization of surface exposed Tie. Shown is an image of fluorescence from Tie_-2xHA_ (magenta), and an overlay image of phase contrast (grey) with fluorescence from Tie-_2xHA_ (magenta). A representative experiment out of 3 independent biological repeats. **E.** Donor strains: WT (SH337: WT/pLS20_cm_), Δ*tie* (SH483: WT/pLS20_cm_-Δ*tie*), and a *tie* complementing strain (SH485: *amyE*::P_IPTG_-*tie*/pLS20_cm_-Δ*tie*) were grown in the presence of IPTG (1 mM), mixed with recipient cells (BDR2637: *sacA::*P*_veg_-mCherry*) in 1:1 ratio, and the formation of MCs in liquid medium was followed using digital wide-field microscopy. Shown is a quantification of the clustering analysis at the indicated time points after mixing. Representative images for time point 30 minutes are presented in Figure S2C. For each mixture, triplicates were analyzed in parallel, and total area of clusters larger than 500 pixels^2^ was calculated. Data is presented as average values and SEM. Statistical significance between WT and each mutant was calculated using two-way ANOVA. For time 30 min, P-values: (ns) not significant ≥ 0.05. P-value for difference between WT/pLS20_cm_-Δ*tie* and Tie-OE/pLS20_cm_-Δ*tie*: (*) ≤ 0.05. A representative experiment out of 3 independent biological repeats.

The link between conjugation, flagella, and MC formation raised the possibility that the flagellum-induced *tie*, located downstream to P_33_ (Figure 3B), previously implicated in mating pair formation^46^, induces MC assembly. Consistent with earlier studies, a donor carrying a plasmid lacking *tie* (pLS20-Δ*tie*) was impaired in conjugation, a property restored by ectopic *tie* overexpression (Figure 5B). Biochemical fractionation of donor cells expressing Tie-_2xHA_ indicated the protein to be localized to the cell wall fraction, consistent with its putative role as an adhesin, and according to its expression level in the various flagellar mutants (Figure 3F and Figure 5C). Moreover, immunofluorescence microscopy exposed the presence of distinct Tie-_2xHA_ foci, decorating the bacterial surface (Figure 5D), supporting its involvement in MC development. We thus assessed Tie impact on MC formation using quantitative clustering analysis. The clustering efficiency of a *tie* deficient donor (pLS20-Δ*tie*) was notably reduced (Figure 5E and Figure S2C), while ectopic expression of *tie* significantly enhanced MC formation in the presence of pLS20, suggesting Tie contribution to this process.

In sum, these findings indicate that flagella induce the expression of conjugation genes in a small subpopulation, demarcated by P_33_ activation, which coincides with MC assembly, and hence conjugation.

### Sensing flagella rotation is required for the expression of conjugation genes

The flagellum harbors a secretory system, which is a key element in the intrinsic circuits hardwired to the process of flagella biogenesis (Figure S3A)^49^. Testing if mutants within these circuits expand their scope to regulate conjugation revealed no significant impact on the process (Figure S3A-S3B). Furthermore, conjugation of Δ*hag* donor could not be rescued in trans by the presence of a hyper-conjugative donor overexpressing RapA (Figure 6A-6B), excluding the involvement of a secreted factor and highlighting that flagella are required strictly in cis.

**Figure 6:**
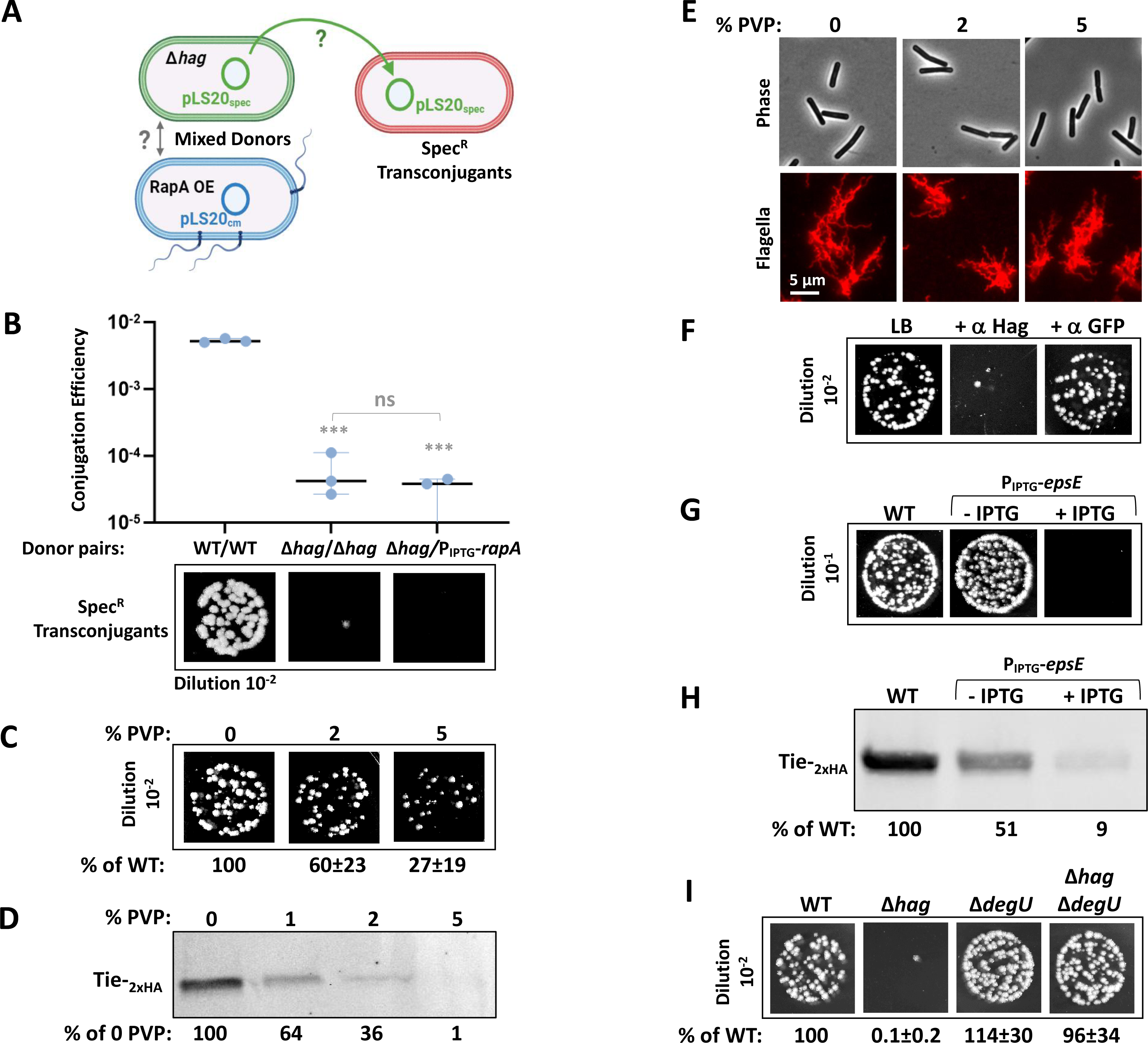
Flagella rotation dictates pLS20 conjugation. **A.** Schematics describing the experimental design to test the ability of a donor strain harboring pLS20_cm_ and over-expressing *rapA* (RapA OE) to suppress in trans the conjugation defect of a Δ*hag* donor strain harboring pLS20_spec_. Conjugation was assayed by monitoring spectinomycin resistant transconjugants. **B.** Corresponds to the experimental design described in (A). Pairs of donor strains: WT (SH337: WT/pLS20_Cm_) with WT (SH342: WT/pLS20_spec_), Δ*hag* (SH443: Δ*hag*/pLS20_Cm_) with Δ*hag* (MBS17: Δ*hag*/pLS20_spec_), and Δ*hag* (MBS17: Δ*hag*/pLS20_spec_) with P_IPTG_-*rapA* (SH450: *amyE*::P_IPTG_-*rapA*/pLS20_Cm_), were grown in the presence of IPTG (0.1 mM), mixed with recipient cells (SH345: *sacA::kan*) in 1:1:1 ratio, and incubated for 20 minutes. Serial dilutions were then spotted either on LB agar containing spectinomycin and kanamycin or solely kanamycin, to specifically select for pLS20_spec_ derived transconjugants and recipients, respectively. Upper panel: Conjugation efficiencies calculated as number of pLS20_spec_-derived transconjugants/number of recipients. At least 3 independent biological repeats were conducted. Data shown as scatter dot plot with a line indicating the median with range. Statistical significance between WT/WT and indicated donor pairs was calculated using one-way ANOVA. P values: (ns) not significant ≥ 0.05, (***) ≤ 0.0005. Lower panel: Representative images of spotted conjugation mixtures (10^-2^ dilution) over LB agar containing spectinomycin and kanamycin, selecting for pLS20_spec_ transconjugants. **C.** Donor (SH337: WT/pLS20_cm_) and recipient (SH345: *sacA::kan*) cells were grown in LB medium supplemented with indicated concentrations of polyvinylpyrrolidone 360 (PVP), mixed in 1:1 ratio, incubated for 20 minutes, and serial dilutions were spotted either on LB agar containing chloramphenicol and kanamycin or solely kanamycin, selecting for transconjugants and recipients, respectively. Shown are images of spotted conjugation mixtures (10^-2^ dilution) over LB agar containing chloramphenicol and kanamycin, selecting for transconjugants. Indicated conjugation efficiencies were calculated as % of WT conjugation efficiency. Presented are average values and SD of at least 3 independent biological repeats. **D.** Whole cell lysates were extracted from WT donor cells (SH461: WT/pLS20_cm_-*tie_-_*_2xHA_) grown in LB medium supplemented with indicated concentrations of PVP and subjected to western blot analysis using anti-HA antibodies. Corresponding stain-free total protein analysis is presented for comparison in Figure S3C. Tie_-2xHA_ expression levels, normalized to stain-free total protein, were calculated as % of 0 PVP. Shown is a representative experiment out of 3 independent biological repeats. **E.** Cells harboring modified flagellin *hag*^T209C^ (DS1895: *amyE*::P*_hag_-hag*^T209C^) were grown in LB medium supplemented with indicated concentrations of PVP, flagellin was stained with Alexa Fluor 594 C_5_ maleimide, and cells were visualized by fluorescence microscopy. Shown are images of phase contrast (grey) (upper panels) and corresponding stained flagella (red) (lower panels). **F.** Donor cells (SH337: WT/pLS20_cm_) were grown in LB medium supplemented with anti-Hag or anti-GFP antibodies as indicated, mixed with recipient cells (SH345: *sacA::kan*) in 1:1 ratio, and incubated for 20 minutes. Shown are images of spotted conjugation mixtures (10^-2^ dilution) over LB agar containing chloramphenicol and kanamycin, selecting for transconjugants. Representative images out of 3 independent biological repeats. **G.** Donor strains: WT (SH337: WT/pLS20_cm_) and P_IPTG_-*epsE* (LZ53: *amyE*::P_IPTG_-*epsE*/pLS20_cm_) were grown in the absence or presence of IPTG (0.5 mM) and mixed with recipient cells (SH345: *sacA::kan*) in 1:1 ratio, and incubated for 20 minutes. Shown are images of spotted conjugation mixtures (10^-1^ dilution) over LB agar containing chloramphenicol and kanamycin, selecting for transconjugants. Representative images out of 3 independent biological repeats. **H.** Whole cell lysates were extracted from donor strains: WT (SH461: WT/ pLS20_cm_-*tie_-_*_2xHA_) and P_IPTG_-*epsE* (SH582: *amyE*::P_IPTG_-*epsE*/pLS20_cm_-*tie_-_*_2xHA_) grown in the absence or presence of IPTG (1 mM), and subjected to western blot analysis using anti-HA antibodies. Stain-free total protein analysis is presented for comparison in Figure S3D. Tie_-2xHA_ expression levels, normalized to stain-free total protein, were calculated as % of WT. Shown is a representative experiment out of 3 independent biological repeats. **I.** Donor strains: WT (SH337), Δ*hag* (SH443), Δ*degU* (SH592), and Δ*hag* Δ*degU* (SH593) harboring pLS20_cm_, were mixed with recipient cells (SH360: *sacA::spec*) in 1:1 ratio, incubated for 20 minutes, and serial dilutions were spotted either on LB agar containing chloramphenicol and spectinomycin or solely spectinomycin, selecting for transconjugants and recipients, respectively. Shown are images of spotted conjugation mixtures (10^-2^ dilution) over LB agar containing chloramphenicol and spectinomycin, selecting for transconjugants. Indicated conjugation efficiencies were calculated as % of WT conjugation efficiency. Presented as average values and SD of at least 3 independent biological repeats.

The finding that the flagellum paralyzed Δ*motA* mutant showed a strong conjugation deficiency that was associated with reduced conjugation gene expression (Figure 2C-2E and Figure 3D-3F), raised the possibility that flagellar rotation might be the signal for activation of the conjugation genes. To address this possibility, we grew the bacteria in media containing polyvinylpyrrolidone (PVP), to increase viscosity, and hence the load imposed on the flagella. Indeed, increased viscosity led to a reduction in both conjugation efficiency and Tie-_2xHA_ expression (Figure 6C-6D and Figure S3C), despite the cells retaining intact flagella (Figure 6E). Furthermore, a strong reduction in conjugation was evident upon escalating the load on the flagella without affecting viscosity, by growing the bacteria in the presence of anti-Hag antibodies, which crosslink the flagellar filaments (Figure 6F).

To further establish the linkage between flagella rotation and conjugation, we ectopically overexpressed *epsE*, encoding a flagellar clutch, which impedes flagellar rotation^50^, in donor cells. Remarkably, EpsE induction caused a dramatic reduction in Tie-_2xHA_ expression that was associated with a severe conjugation blockage (Figure 6G-6H and Figure S3D). As the DegS-DegU two-component system was implicated in relaying flagella rotation to gene expression^51,52^, the potential involvement of this mechanosensing pathway in conjugation was examined. Conjugation efficiency of a strain lacking the response regulator DegU was similar to that of WT (Figure 6I). Nevertheless, the conjugation defect of the Δ*hag* strain was fully suppressed upon *degU* deletion (Figure 6I), indicating an inhibitory effect of DegU transcriptional factor on conjugation gene expression.

Cumulatively, our results show that upon sensing rotating flagella, a signal is generated to alleviate the inhibitory effect of DegU on conjugation, thereby tying motility with conjugation. Evolution of such plasmid-host molecular cross-talk likely reflects a strategy to activate conjugation when spreading into new habitats is within reach.

### Conjugation-motility coupling fosters spreading into new ecological niches

To demonstrate the prospect that conjugation by flagellated donors is advantageous over non-flagellated ones in promoting invasion into new habitats, we exploited the *degU* mutation, which uncouples motility from conjugation gene expression. Accordingly, we compared the spreading efficiency of transconjugants derived from motile (Δ*degU*) and non-motile (Δ*degU* Δ*hag*) donors to a restrictive niche. We simulated an ecological system composed of two niches, one antibiotic- free while the other supplemented with chloramphenicol and spectinomycin, selectively permitting the growth of transconjugants (Figure 7A). The motile and non-motile donors were spotted over the antibiotic-free zone at equal distances from a motile GFP expressing recipient, and the expansion of transconjugants into the selective region was monitored (Figure 7A). Flagellated donors rapidly encountered the recipients, and transconjugants emerging from the donor-recipient intersection expanded over time into the restrictive zone (Figure 7B and Figure S4). In contrast, despite similar conjugation proficiency (Figure 6I), transconjugants derived from the non-motile donor failed to invade the selective zone even after 28 hours of incubation (Figure 7B and Figure S4). As a control, no expansion into the antibiotic-containing area was seen when monitoring identical strains lacking pLS20 (Figure 7B). These results insinuate that being motile is beneficial for reaching remotely located recipients and spreading into new niches, highlighting the evolutionary pressure to tie conjugation with flagella rotation.

**Figure 7:**
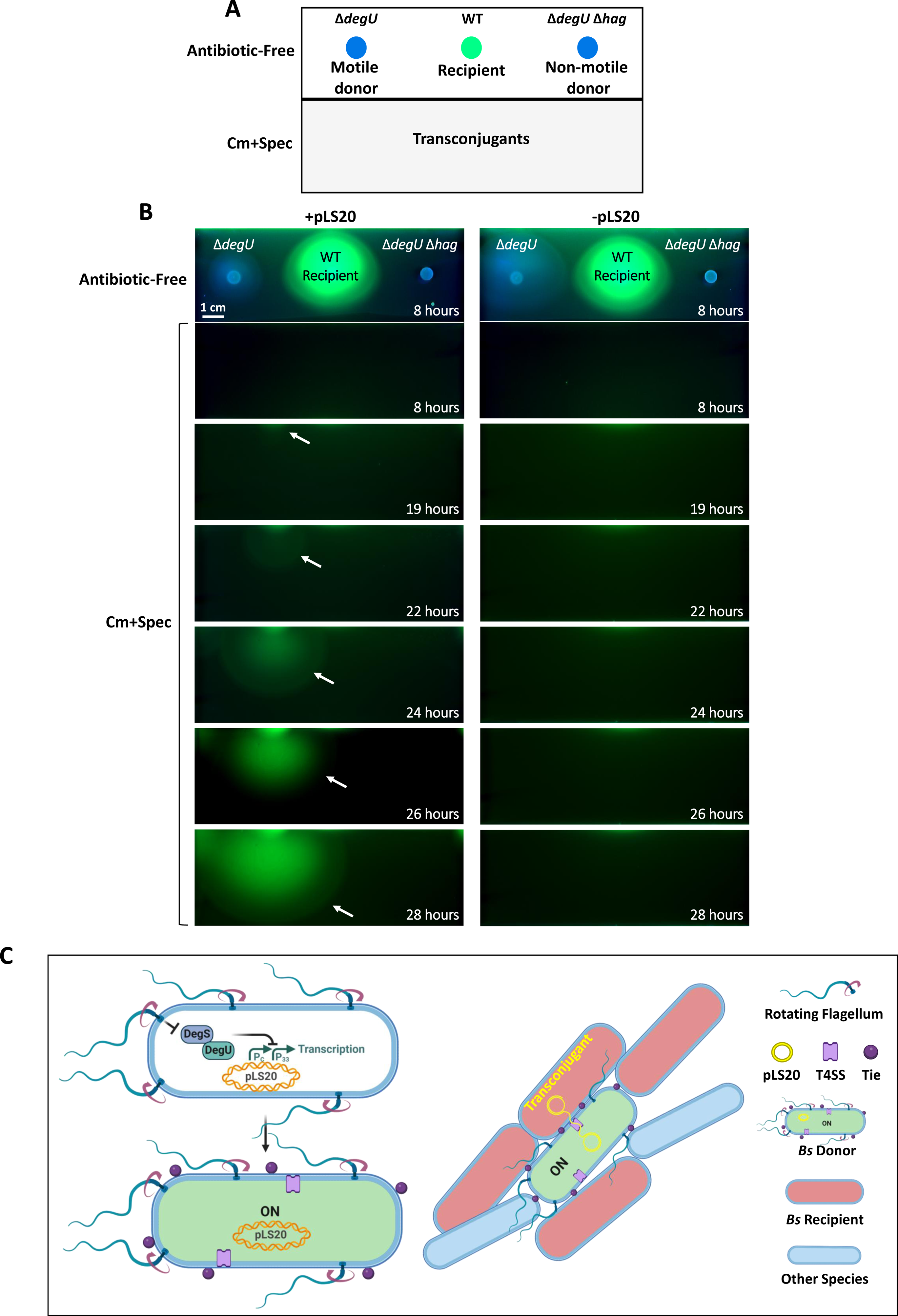
The flagella-conjugation link allows spreading into new ecological niches. **A.** A diagram outlining the experimental design to demonstrate the impact of flagella on pLS20_cm_ spread. A split-plate soft agar assay was devised, wherein the top half of a rectangular plate contained antibiotic-free LB and the bottom half contained LB supplemented with chloramphenicol and spectinomycin (Cm+Spec). The entire plate was layered with soft agar, and motile (Δ*degU*) and non-motile (Δ*hag* Δ*degU*) donors (blue circles) were spotted at equal distances from a WT motile GFP expressing recipient (green circle) on the antibiotic-free zone. The soft agar allows bacterial growth, swimming motility and conjugation. The emergence of motile GFP-expressing transconjugants resistant to chloramphenicol and spectinomycin on the antibiotic selective region could then be monitored over time. **B.** Corresponds to the experimental design in (A). Bacteria were grown to mid-logarithmic phase and spotted on the antibiotic-free region of the plate. Left panel: plates were spotted with Δ*degU* (SH592: Δ*degU*/pLS20_cm_), Δ*hag* Δ*degU* (SH593: Δ*hag* Δ*degU*/pLS20_cm_), and WT (AR16: P*_rrnE_*-*gfp*) strains. Right panel: plates were spotted with Δ*degU* (SH590: Δ*degU*), Δ*hag* Δ*degU* (SH609: Δ*hag* Δ*degU*), and WT (AR16: P*_rrnE_*-*gfp*) strains. Presented are overlay images of colorimetric channel (blue) with fluorescence from GFP (green), captured at the indicated time points post incubation using ChemiDoc MP imaging system. Upper panels: images from the antibiotic-free region after 8 hours of incubation. Lower panels: a series of images from the antibiotic-containing region (Cm+Spec), captured at the indicated time points post incubation. White arrows point at the emergence and spread of transconjugants. Images of the entire plates (left panels) are presented in Figure S4. Shown is a representative experiment out of 3 independent biological repeats. **C.** A model depicting donor flagella-driven activation of conjugation gene expression and MC formation in liquid milieu. Left Panel: schematic illustrating a motile *B. subtilis* (*Bs*) donor cell, in which flagellar rotation leads to alleviation of the inhibitory effect of DegS-DegU two component system on pLS20. Consequently, transcription of crucial conjugation genes, including the T4SS machinery and the adhesin Tie, regulated by P_C_ and P_33_ are activated, thereby switching to a conjugation “ON” state. Right Panel: Assembly of MC, composed of *B. subtilis* and other species, induced by a conjugation primed “ON” cell, providing a stable platform for mating pair formation and plasmid DNA transfer.

## Discussion

To optimize horizontal distribution, conjugative elements must sense host physiological cues to activate conjugation gene expression under favorable conditions. Here we discovered that pLS20, a prevalent conjugative plasmid found across *Bacillus* species^18^, has evolved to plug-in to the host mechanosensing signal transduction pathway, designed to discern flagella rotation. This adaptation enables the plasmid to coordinate its spread with the motile phase of the bacterial life cycle, which typically coincides with the bacterial necessity to relocate from challenging microenvironments^e.g.53^. Furthermore, we demonstrate that this linkage accelerates dissemination of antibiotic resistance through conjugation. Upon detecting propelled flagella, pLS20 conjugative genes, including the T4SS machinery, are induced in a subset of cells, becoming primed to exploit potential conjugation opportunities. As conjugation demands a fastened donor-recipient pairing to allow DNA delivery, pLS20 orchestrates clustering of the donor with recipient bacteria, aided by the plasmid-encoded Tie protein. These MCs facilitate intimate proximity between donor and recipient bacteria, providing a hub for mating pair formation (Figure 7C). Interestingly, foreign species could be efficiently recruited to *B. subtilis* donor- derived MCs, alluding to the potential of these dynamic cellular assemblages to drive genetic exchange among species within natural niches (Figure 7C).

Most well-studied conjugation systems favor solid surfaces, offering stability and high cell density, crucial for mating pair establishment and plasmid delivery^4^. In fact, biofilm cellular assemblages emerge as significant hotspots for horizontal gene transfer among bacteria residing in natural habitats^e.g.54,55^, a phenomenon that can even be promoted by artificial bacterial adhesion^56^. Consistently, some conjugative elements were found to actively temper host motility, possibly facilitating the establishment of cell-to-cell connection. A prominent example is the enterobacteria conjugative plasmid R27, encoding factors that downregulate flagella synthesis^57^. Surprisingly, pLS20 has evolved to synchronize activation of conjugation with functional flagella, although *B. subtilis* cells are known to form robust biofilms^e.g.28,58,59^. This feature provides the plasmid with the benefit of spreading into remote destinations, seizing opportunities to invade new hosts of the same or diverse species. The strategy of inducing the formation of temporal MCs can be advantageous over biofilms, which confine donor bacteria to relatively predetermined, fixed niches.

The capacity of mobile genetic elements to be tuned with the host physiological state is a prevalent trait that appears to maximize their fitness. Numerous prophages, such as lambda, trigger the lysogeny-to-lysis transition in response to host DNA damage, a synchronization facilitated through host SOS regulatory factors^60,61^. Likewise, the excision of the *B. subtilis* mobile genetic element, ICEBs1, was shown to be induced by the global DNA-damage SOS response^62^, whereas the *tra* operon of the F-family conjugative plasmid is governed by the host aerobic respiration control factor ArcA, hence directly linking conjugation with oxygen levels^63^. Importantly, conjugative elements can also interfere with host physiology by modifying global gene expression, and may even delay entry into biofilm or sporulation developmental states^64–66^. Similarly, transcriptome analysis of *B. subtilis* cells harboring pLS20, revealed a substantial impact on host gene expression, including genes involved in metabolic pathways, cell envelope, motility, signal transduction and regulatory pathways^66^. Thus, mobile genetic elements can sense and intervene with host physiology.

An important facet of conjugative plasmids is their maintenance in a default ‘OFF’ state, presumably to avoid the fitness cost that would be associated with their constitutive expression^67,68^. Regulation of pLS20 has been primarily attributed to the modulation of the P_C_ promoter activity^24,48^. However, the discovery of additional promoters, including P_33_ in this study, along with transcription terminators within the conjugation operon of pLS20^47,69,70^, highlight the presence of additional layers of regulation facilitating a switch from conjugation ‘OFF’ to ‘ON’ state. Fascinatingly, pLS20 senses the host flagellar rotation that serves as a mechanotransmitter to activate P_C_ and P_33_ promoters in a subset of cells, via the DegS-DegU system. Interestingly, *B. subtilis* flagella has been implicated as a mechanotransmitter, dictating developmental processes such as competence and biofilm formation^51,52,71^. Bacterial mechanosensing appears as a widespread strategy for integrating environmental cues to coordinate complex physiological responses^e.g.72,73^. Mechanosensing properties have also been attributed to type IV pili in *Pseudomonas aeruginosa* in detecting solid surfaces and host cells to activate virulence pathways^73–76^, whereas in *Caulobacter crescentus*, surface sensing can be mediated both via the tight adherence pilus and flagella^77,78^.

Flagella have also been implicated in serving as an adhesin in several bacteria, including pathogenic *E. coli*, *P. aeruginosa*, *Salmonella* Typhimurium, *Clostridium difficile*, mediating contact with host cells or even with abiotic surfaces^e.g.42,43,79–83.^ Further, flagella of the marine bacterium *P. marina* have been demonstrated to promote intraspecies aggregation in artificial sea water^44^. It is conceivable that flagella might mediate an initial contact during pLS20 conjugation, enhancing donor-recipient proximity, aiding Tie to securely fasten bacterial cells to form stable MCs and hence mating pair formation. However, according to our data, flagella contribution to MC is restricted to donor bacteria, suggesting that in this case flagella by itself cannot simply act as an aggregation facilitator.

Our findings unravel the riddle of efficient conjugation of pLS20 in liquid medium despite lacking an apparent pilus, thereby shedding light on novel mechanisms driving horizontal gene transfer in microbial communities residing in fluid environments. The implications of this study could inform strategies for controlling the spread of antibiotic resistance genes through conjugation in Gram-positive bacteria.

## Acknowledgments

We thank members of the Ben-Yehuda and Rosenshine laboratories for valuable discussions. We are grateful to Daniel B. Kearns (Indiana U) for providing bacterial strains and anti-Hag antibodies, and Mitsuhiro Itaya (Keio U), and David Rudner (Harvard U) for providing bacterial strains and plasmids. This work was supported by the ERC Synergy grant (810186) awarded to S. B-Y and I.R., and by the German Research Foundation (DFG) Priority Program SPP2389 awarded to S. B-Y. S.B was funded by the Golda Meir postdoctoral fellowship.

## Methods

### Bacterial strains and plasmids

All *B. subtilis* strains used in this study are derivatives of wild type PY79 strain^1^. *B. cereus* (OS4) *and B. megaterium* (OS2)^2^ are wild type soil isolates (laboratory stock). All bacterial strains and plasmids are described in Table S1. All primers used in this study are listed in Table S2. Plasmid constructions were performed using standard molecular biology methods in *E. coli* DH5α.

### General growth conditions

All general methods for *B. subtilis* were carried out as described previously^3^. *B. subtilis* cultures were inoculated from an overnight culture and growth was carried out to the desired OD_600_ at 37°C in LB medium (Difco). For strains harboring genes under inducible promoters, 0.1-1 mM Isopropyl-β-D-thiogalactopyranoside (IPTG, Sigma-Aldrich) was added to the medium, as indicated. Antibiotics were used at the following concentrations: chloramphenicol 5 μg/ml (Sigma-Aldrich), kanamycin 10 μg/ml (US Biological), erythromycin 1 μg/ml (Sigma-Aldrich), lincomycin 25 μg/ml (Sigma-Aldrich), spectinomycin 100 μg/ml (Sigma-Aldrich), tetracycline 10 μg/ml (Sigma-Aldrich). For *E. coli* strains, ampicillin 100 ug/ml (Sigma-Aldrich) was used for selection.

### Conjugation assay by CFU

Conjugation assay in liquid medium was based on a previously described protocol^4^. *B. subtilis* donor cells, harboring pLS20 or its derivatives, and recipient cells, lacking a plasmid, were grown to OD_600_ 0.8 at 37°C in LB medium, mixed in 1:1 ratio and incubated for 20 minutes at 37°C under static conditions to permit conjugation. Serial dilutions were spotted (100 μl) on LB agar containing appropriate antibiotics to specifically select for transconjugants, donors, and recipients. Conjugation efficiencies were calculated as number of Transconjugant CFUs/number of recipient CFUs.

### Conjugation in the presence of PVP

*B. subtilis* donor cells, harboring pLS20, and recipient cells, lacking a plasmid, were grown to OD_600_ 0.8 at 37°C in LB medium supplemented with different PVP (Polyvinylpyrrolidone 360, Sigma-Aldrich) concentrations to increase the viscosity of the medium. Conjugation assay was conducted in the presence of PVP as described above.

### Conjugation in the presence of antibodies

*B. subtilis* donor cells, harboring pLS20 were grown for 2 hours at 37°C in LB medium supplemented with anti-Hag (a gift from Daniel B. Kearns) or anti-GFP antibodies (Invitrogen) at a final concentration of 1:100. In parallel, WT recipient cells, lacking a plasmid, were grown to OD_600_ 0.8 at 37°C in LB medium. Conjugation assay was then conducted as described above.

### Split-plate soft-agar conjugation assay

For conjugation on soft agar, freshly prepared LB containing 0.3% agar was layered over an LB-rectangular plate containing 1.5% agar that was split into two halves. The top half contained antibiotic-free LB, permitting bacterial growth, swimming motility, and conjugation. The bottom part contained LB supplemented with chloramphenicol and spectinomycin antibiotics for selection of transconjugants. The combined plates were composed by excising the LB bottom half, and replacing it with LB supplemented with chloramphenicol and spectinomycin. Strains harboring or lacking pLS20, and recipient cells constitutively expressing GFP, were grown to mid-logarithmic phase in LB and concentrated to OD_600_ 5.0. Cell suspension (3 μl) was spotted over the antibiotic-free region. Plates were incubated at 37°C and imaged over time using ChemiDoc MP imaging system (Bio-Rad).

### Motility assay

Motility assay was based on previously described protocol^5^. Cells were grown to mid-logarithmic phase in LB and concentrated to OD_600_ 5.0. Cell suspension (5 μl) was spotted on freshly prepared LB plates containing 0.3% agar, incubated at 37°C for 7 hours, and imaged using Epson Perfection V800 Photo scanner.

### Protein extraction and western blot analysis

#### Protein extraction

For preparation of whole cell lysates, *B. subtilis* cells were grown to OD_600_ 0.8 at 37°C in LB medium. Cells were harvested and suspended in lysis buffer containing 10 mM Tris-HCl pH 8.0, 10mM MgCl_2_, 0.5 mg/ml Lysozyme, 5 μg/ml DNaseI, and 1x Halt™ protease inhibitor cocktail (Thermo Fisher Scientific), and incubated at 37°C for 10 minutes. Following cell lysis, samples were incubated at 95°C for 10 minutes with 1x Laemmli sample buffer (Bio-Rad).

Cell wall protein extraction was carried out as described previously^6^. Briefly, *B. subtilis* cells were grown to OD_600_ 0.8 at 37°C in LB medium. Cells were harvested, and washed twice with 10 mM Tris-HCl, pH 8.0. Cell pellet was then resuspended in a buffer containing 1.5 M LiCl, 25 mM Tris-HCl, pH 8.0, and kept on ice for 10 minutes. The suspension was then centrifuged, and supernatant collected. Proteins were precipitated using 10% Trichloroacetic acid (TCA) (w/v) at 4°C for one hour. Suspension was centrifuged (15,000 rpm, 30 minutes, 4°C), and the pellet was washed 3 times in ice cold Ethanol (100%), air dried, resuspended in 1x Laemmli sample buffer (Bio-Rad) and incubated at 95°C for 10 minutes.

#### Western blot analysis

Protein samples were separated by 4-20% Mini-PROTEAN TGX Stain-Free Precast Gels (Bio-rad), and electroblotted onto a Nitrocellulose membrane with Trans-Blot Turbo Transfer System (Bio-rad). The membrane was blocked for 1 h with 5% skim milk in TBST (50 mM Tris-Cl, pH 7.5, 150 mM NaCl, 1% Tween-20). Blots were then probed with Rabbit anti-HA antibodies (1:5000 in TBST, Invitrogen) followed by Goat anti-Rabbit fluorophore-conjugated secondary antibody Star Bright B700 (Bio-Rad) (1:5,000 in TBST). Gels and blots were imaged using ChemiDoc MP imaging system (Bio-Rad). Stain-Free image was used as total protein loading control.

### Quantitative reverse transcriptase PCR

RNA was extracted from *B. subtilis* cells grown to OD_600_ 0.8 using Direct-zol™ RNA Miniprep plus kit (ZYMO RESEARCH), according to the manufacturer protocol. RNA concentration was determined using NanoDrop™ One (Thermo Scientific). RNA (2 μg) from each sample was treated with TURBO™ DNase (2 Units, Thermo Scientific), and subjected to cDNA synthesis using iScript cDNA synthesis kit (Bio-Rad), according to the manufacturer protocol. qRT-PCR reactions were performed in triplicates using iTaq Universal SYBR Green Supermix (Bio-Rad), and fluorescence detection was performed using Bio-Rad CFX Connect Real time system. RNA expression was normalized to the level of 16S rRNA. To verify that a single product was amplified, a melt curve analysis was performed using the Bio-Rad CFX manager software. The relative gene expression levels were calculated from threshold cycle (C_T_) values using the 2^−ΔΔCT^ method. qRT-PCR primers were designed using Primer3 software (v. 0.4.0, available online).

### Fluorescence microscopy

All fluorescence microscopy imaging was performed using ECLIPSE Ti2 microscope (Nikon, Japan), equipped with Prime BSI camera (Photometrics, Roper Scientific, USA). System control and image analysis were performed using NIS-Elements AR Analysis (version 5.30.07, Nikon, Japan), and FIJI (Image J).

Fluorescence microscopy of conjugation using pLS20-P_IPTG_-*gfp* reporter: *B. subtilis* donor strain constitutively expressing mCherry and LacI, and harboring pLS20-P_IPTG_-*gfp*, was grown to OD_600_ 0.8 at 37°C in LB medium, and mixed in 1:1 ratio with recipient cells grown under similar conditions. The mixture was incubated for 90 minutes at 37°C under static conditions to permit conjugation and expression of GFP in the recipient cells. At indicated time points, samples were treated with 2% paraformaldehyde for 10 minutes at 25°C, washed with 1xPBS (Phosphate Buffered Saline, pH7.4), spotted on microscope slides (Marienfeld), covered with poly-L-lysine coated coverslips, and visualized by fluorescence microscopy. Recipient cells, expressing GFP, were considered transconjugants.

Fluorescence microscopy of conjugation using pLS20-*ssb*-*yfp* reporter: *B. subtilis* donor strains harboring pLS20-*ssb-yfp*, and recipient cells constitutively expressing mCherry, were grown to OD_600_ 0.8 at 37°C in LB medium, mixed in 1:1 ratio and incubated for 60-80 minutes at 37°C under static conditions to permit conjugation and expression of SSB-YFP in the recipient cells. At indicated time points, samples were treated with 2% paraformaldehyde for 10 minutes at 25°C, washed with 1xPBS (Phosphate Buffered Saline, pH7.4), spotted on microscope slides (Marienfeld), covered with poly-L-lysine coated coverslips, and visualized by fluorescence microscopy. Recipient cells, expressing mCherry and displaying SSB-YFP foci, were considered transconjugants. For time lapse microscopy analysis, donor and recipient strains were grown to OD_600_ 0.8 at 37°C in LB medium, mixed in 1:1 ratio, and the mixture was mounted onto a metal ring (A-7816, Invitrogen) filled with LB agarose (0.6%). Cells were incubated in a temperature and humidity-controlled chamber (Okolab) at 37°C, and visualized over time by fluorescence microscopy.

#### Evaluating conjugative promoter activity

*B. subtilis* strains harboring P_C_-*gfp* or P_33_-*gfp* transcriptional reporters were grown to OD_600_ 0.8 at 37°C in LB medium, harvested, washed with 1xPBS, spotted on microscope slides (Marienfeld), covered with poly-L-lysine coated coverslips and imaged by fluorescence microscopy. GFP levels for each cell were determined

#### Visualizing MC formation

*B. subtilis* donor cells, harboring pLS20-*ssb-yfp* in the presence or absence of the P_33_-*gfp* transcriptional reporter, and recipient cells constitutively expressing mCherry, were grown to OD_600_ 0.8 at 37°C in LB medium and mixed in 1:1 ratio. The mixed cells were mounted onto a metal ring (A-7816, Invitrogen) filled with LB agarose (0.6%), incubated in a temperature and humidity-controlled chamber (Okolab) at 37°C for 60-80 minutes and visualized by fluorescence microscopy.

#### Visualizing multi-species MC formation

*B. subtilis* donor cells, harboring pLS20, *B. subtilis* recipient cells constitutively expressing mCherry, and *B. megaterium* or *B. cereus* strains, were grown to OD_600_ 0.8 at 37°C in LB medium and mixed in 1:1:1 ratio. The mixed cells were mounted onto a metal ring (A-7816, Invitrogen) filled with LB agarose (0.6%), incubated in a temperature and humidity-controlled chamber (Okolab) at 37°C for 45 minutes and visualized by fluorescence microscopy.

#### Flagella staining and visualization

Flagella visualization was carried out as previously described^7^, with modifications. *B. subtilis* cells expressing modified flagellin (*amyE*::P*_hag_- hag*^T209C^) were harvested at OD_600_ 0.8, washed with 1xT-BASE buffer [15 mM (NH_4_)_2_SO_4_, 80 mM K_2_HPO_4_, 44 mM KH_2_PO_4_, 3.4 mM sodium citrate, and 3 mM MgSO_4_] (pH 8), resuspended in 1xT-BASE buffer (pH 8) containing 5 μg/ml Alexa Fluor 594 C_5_ maleimide (Molecular Probes, Thermo Fisher Scientific), and incubated for 5 minutes at room temperature. Cells were then washed with 1xT-BASE buffer (pH 8), mounted onto a metal ring (A-7816, Invitrogen) filled with LB agarose (1.5 %) and visualized by fluorescence microscopy.

### Immunofluorescence

*B. subtilis* cells grown to OD_600_ 0.8 at 37°C in LB medium were harvested and washed once with 1xPBS. Cells were then treated with Rabbit anti-HA primary antibody (1:200, Invitrogen) for 40 minutes at room temperature, washed two times with 1xPBS, and subsequently treated with Alexa 647 conjugated secondary antibody (1:2000, Abcam) for 30 minutes at room temperature. Cells were then washed two times with 1xPBS, spotted on microscope slides (Marienfeld), covered with poly-L-lysine coated coverslips and visualized by fluorescence microscopy.

### MC formation analysis by digital microscopy

*B. subtilis* donor strains, harboring pLS20, and recipient cells, constitutively expressing mCherry, were grown to OD_600_ 0.8 at 37°C in LB medium, mixed in 1:1 ratio, and mixtures were placed in a 96 well BLACK, CELLSTAR® plate (Greiner). Clustering of red recipient cells was followed over time at 37°C, under static conditions, in digital wide-field microscopy (BioTek Cytation 5, Agilent). Automated Image capturing was performed at 10 minute intervals using a 10X objective and the BioTek Gen5 Software. Image processing was performed with Fiji-ImageJ. To estimate cluster formation, images of mCherry channel were subjected to thresholding and analyzed for particles larger than 500 pixels^2^.

### Monitoring conjugative promoter activity

*B. subtilis* cells harboring P_C_-*gfp* or P_33_-*gfp* transcriptional reporters were grown to OD_600_ 0.8 at 37°C in LB medium, harvested, washed and resuspended in 1xPBS. <u>Fluorescence of spotted cultures:</u> Samples (10 μl) were spotted in triplicates on LB agar plates and incubated at 37°C for 18 hours. Plates were imaged using ChemiDoc MP imaging system (Bio-Rad).

#### Plate reader

Samples were tested in triplicates in a 96 well BLACK, CELLSTAR® plate (Greiner). Fluorescence intensity and OD_600nm_ were measured by Spark 10M (Tecan) multiwell fluorometer plate reader. GFP intensities were normalized to the OD_600_ values following the subtraction of the background GFP intensity of a WT donor strain (SH337, pLS20) lacking GFP.

### Statistical analysis

Statistical analysis was performed using GraphPad Prism software. P values obtained are indicated in the respective figures.

## Supplemental Figure Legends

**Figure S1:**
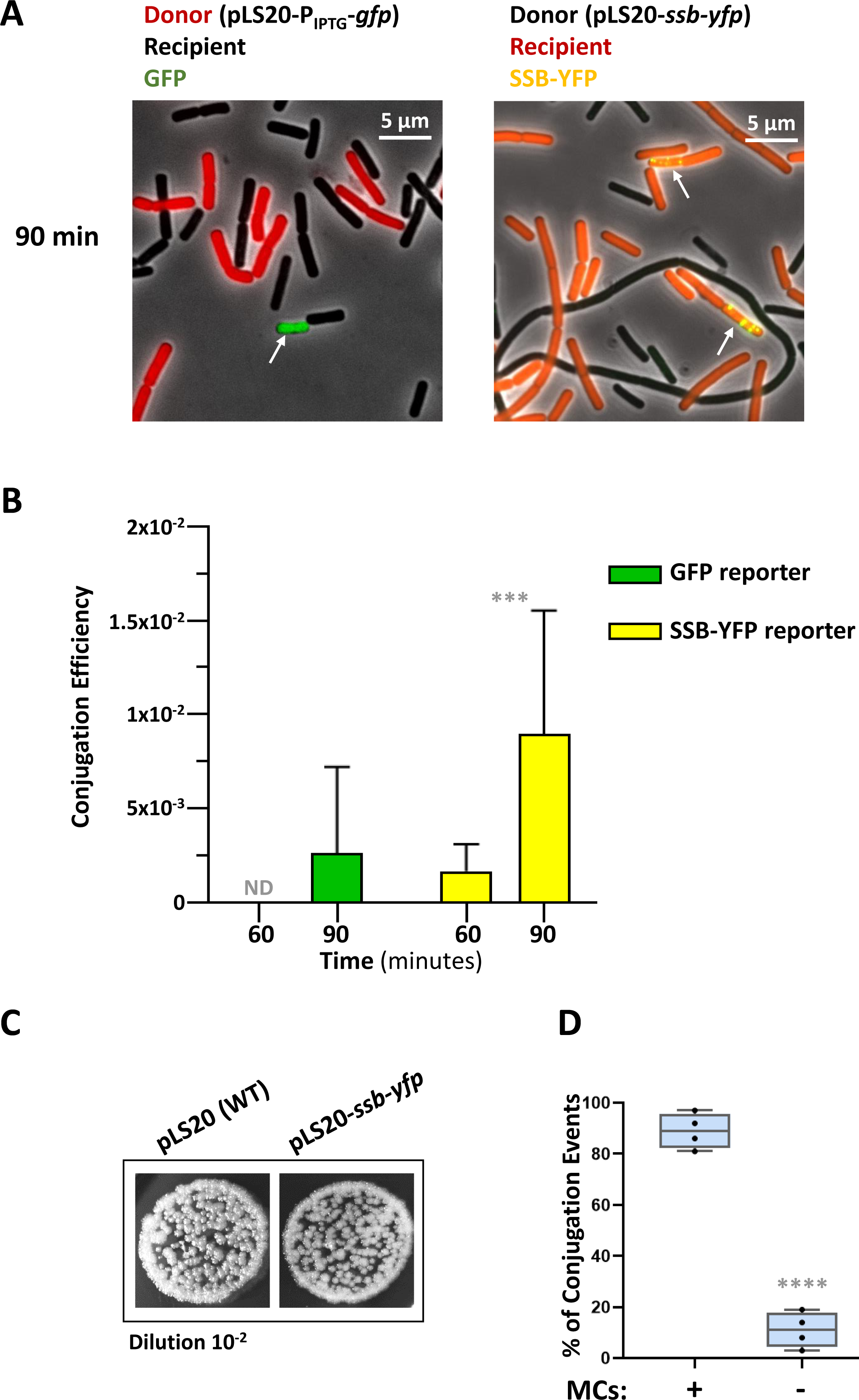
Developing a system for visualizing pLS20 conjugation events. **A.** Left panel: Donor cells (red) (SH363: WT, *sacA::*P*_veg_-mCherry, amyE::lacI*/pLS20_spec_-P_IPTG_*-gfp*) were mixed with recipient cells (dark) (PY79:WT), incubated for 90 minutes and visualized by fluorescence microscopy. Shown is an overlay image of phase contrast (grey) with fluorescence from mCherry (red) and GFP (green). GFP synthesis from pLS20_spec_-P_IPTG_*-gfp* is repressed in donor bacteria by LacI that is lacking from recipient cells. Thus, recipient cells, expressing GFP, were scored as transconjugants. Right panel: Donor cells (dark) (SH347: WT/pLS20_cm_-*ssb-yfp*) were mixed with recipient cells (red) (BDR2637: *sacA::*P*_veg_-mCherry*) in 1:1 ratio, incubated for 90 minutes and visualized by fluorescence microscopy. Shown is an overlay image of phase contrast (grey), with fluorescence from mCherry (red) and SSB-YFP (yellow). Recipient cells, expressing mCherry and displaying SSB-YFP foci, were scored as transconjugants. Arrows highlight transconjugant cells. Representative images out of 3 independent biological repeats. **B.** Conjugation efficiencies of the strains described in (A) were calculated as the number of transconjugants/number of recipients at the indicated time points. Data is presented as average values and SEM of at least seven fields from a representative experiment out of 3 independent biological repeats. Statistical significance between GFP reporter and SSB-YFP reporter was calculated using two-way ANOVA. P-values: (***) ≤ 0.001. ND-not detected. **C.** Donor strains: (SH337: WT/pLS20_cm_) and (SH347: WT/pLS20_cm_-*ssb-yfp*) were mixed with recipient cells (SH345: *sacA::kan*) in 1:1 ratio, and incubated for 20 minutes. Shown are images of spotted conjugation mixtures (10^-2^ dilution) over LB agar containing chloramphenicol and kanamycin, selecting for transconjugants. Representative images out of 3 independent biological repeats. **D.** Donor cells (SH347: WT/pLS20_cm_-*ssb-yfp*) were mixed with recipient cells (BDR2637: *sacA::*P*_veg_-mCherry*) in 1:1 ratio, placed over semi-solid (0.6%) LB agarose pads, incubated for 80 minutes, and visualized by fluorescence microscopy. Conjugation events demonstrated by transconjugant cells expressing SSB-YFP were quantified and categorized based on their association with MCs. At least 4 independent biological repeats were conducted. Data are shown as box plot graphs. The box is determined by the 25^th^ and 75^th^ percentiles, and whiskers are determined by min and max; the line in the box indicates the median. Statistical significance was calculated using paired t-tests. P-value: (****) ≤ 0.0001.

**Figure S2:**
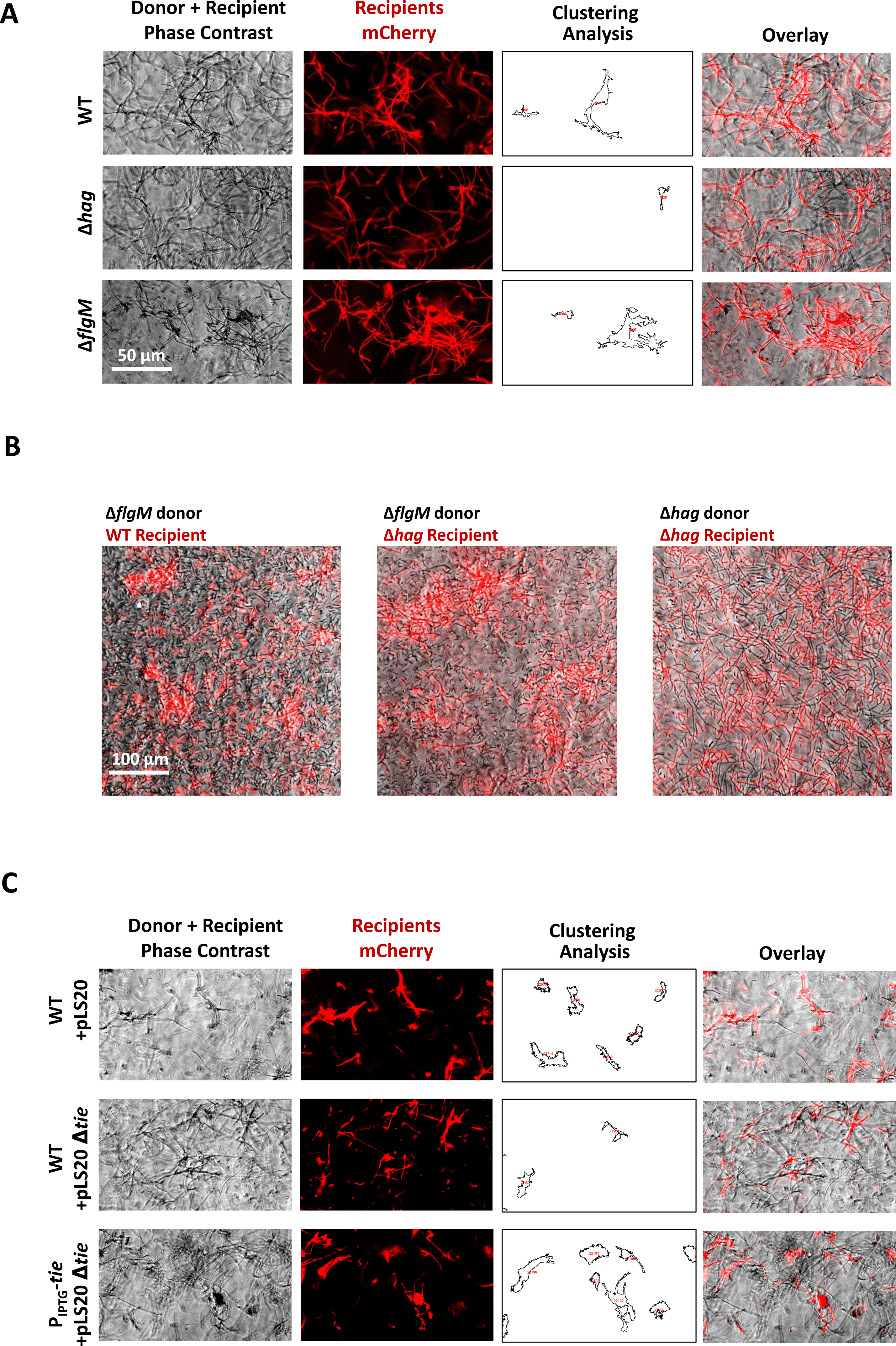
MCs are pLS20 and flagella dependent. **A.** Donor strains (dark) WT (SH337), Δ*hag* (SH443), or Δ*flgM* (SH496) harboring pLS20_cm_, were mixed with recipient cells (red) (BDR2637: *sacA::*P*_veg_-mCherry*) in 1:1 ratio, and the formation of MCs in liquid medium was followed using digital wide-field microscopy. Shown are representative images captured at time point 30 minutes after mixing: phase contrast (grey), fluorescence from mCherry-labeled recipients (red), computed clustering analysis (outlines), and overlay of phase contrast and mCherry fluorescence. Quantification of this experiment is shown in Figure 1F. A representative experiment out of 3 independent biological repeats. **B.** Donor strains (dark) Δ*flgM* (SH496) or Δ*hag* (SH443), harboring pLS20_cm_, were mixed with recipient cells (red) WT (BDR2637: *sacA::*P*_veg_-mCherry*) or Δ*hag* (SH568: Δ*hag, sacA::*P*_veg_-mCherry*) in 1:1 ratio, and the formation of MCs in liquid medium was followed using digital wide-field microscopy. Shown are representative overlay images of phase contrast (grey) and fluorescence from mCherry-labeled recipients (red) captured 30 minutes after mixing. A representative experiment out of 3 independent biological repeats. **C.** Donor strains (dark): WT (SH337: pLS20_cm_), Δ*tie* (SH483: pLS20_cm_-Δ*tie*), and a *tie* complementing strain (SH485: *amyE*::P_IPTG_-*tie*/pLS20_cm_-Δ*tie*) were grown in the presence of IPTG (1 mM) and mixed with recipient cells (red) (BDR2637: *sacA::*P*_veg_-mCherry*) in 1:1 ratio, and the formation of MCs in liquid medium was followed using digital wide-field microscopy. Shown are representative images captured at time point 30 minutes after mixing: phase contrast (grey), fluorescence from mCherry-labeled recipients (red), computed clustering analysis (outlines), and overlay of phase contrast and mCherry fluorescence. Quantification of this experiment is shown in Figure 5E. A representative experiment out of 3 independent biological repeats.

**Figure S3:**
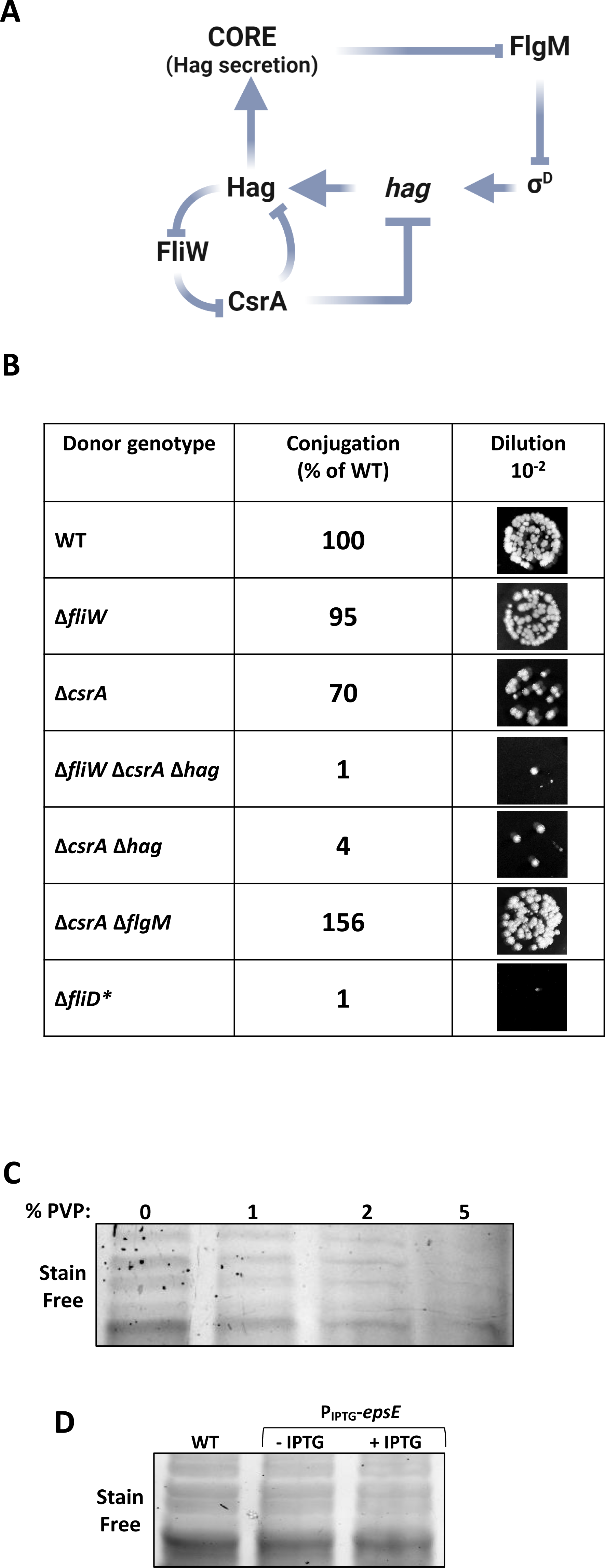
Exploring the mechanism of flagella-mediated conjugation. **A.** Schematics depicting the regulatory circuits in *B. subtilis* cells governing flagella biosynthesis via molecular regulation of *hag* at the transcriptional and translational levels. Arrows highlight activation and T-bars indicate inhibition. Adapted from (Oshiro *et al.*, 2019)^49^. **B.** Donor strains: WT (SH337), Δ*fliW* (GV289), Δ*csrA* (GV290), Δ*fliW* Δ*csrA* Δ*hag* (GV291), Δ*csrA* Δ*hag* (GV293), Δ*csrA* Δ*flgM* (SH381), and Δ*fliD* (SH422), harboring pLS20_cm_ were mixed with recipient cells (SH345: *sacA::kan*) in 1:1 ratio, incubated for 20 minutes, and serial dilutions were spotted either on LB agar containing chloramphenicol and kanamycin or solely kanamycin, selecting for transconjugants and recipients, respectively. Shown are images of spotted conjugation mixtures (10^-2^ dilution) over LB agar containing chloramphenicol and kanamycin, selecting for transconjugants. Indicated conjugation efficiencies were calculated as % of WT conjugation efficiency. A representative experiment out of 3 independent biological repeats. \**fliD* encodes a homolog of flagellar filament cap protein, serving as an extra-cytoplasmic chaperone for polymerization of Hag. **C.** Whole cell lysates were extracted from WT donor cells (SH461: WT/pLS20_cm_-*tie_-_*_2xHA_), grown in LB medium supplemented with different concentrations of PVP. Samples were subjected to SDS-PAGE and stain-free total protein analysis. The corresponding western blot analysis using anti-HA antibodies are presented in Figure 6D. **D.** Whole cell lysates were extracted from donor strains WT (SH461: WT/ pLS20_cm_-*tie_-_*_2xHA_) and P_IPTG_-*epsE* (SH582: *amyE*::P_IPTG_-*epsE*/pLS20_cm_-*tie_-_*_2xHA_), grown in the absence or presence of IPTG (1 mM). Samples were subjected to SDS-PAGE and stain-free total protein analysis. The corresponding western blot analysis using anti-HA antibodies are presented in Figure 6H.

**Figure S4:**
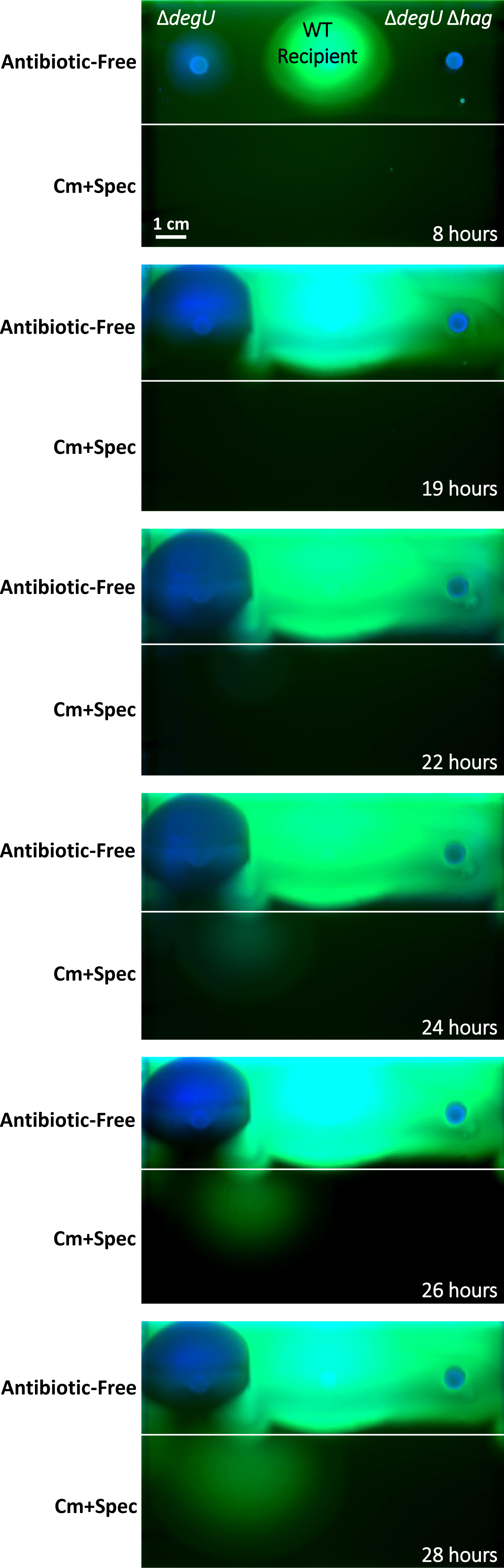
Coupling motility and conjugation is beneficial for invading new niches. Images of entire plates corresponding to the series presented in Figure 7B (left panels). Bacteria were grown to mid-logarithmic phase and spotted on the antibiotic-free region of the plate. Plates were spotted with Δ*degU* (SH592: Δ*degU*/pLS20_cm_), Δ*hag* Δ*degU* (SH593: Δ*hag* Δ*degU*/pLS20_cm_), and WT (AR16: P*_rrnE_*-*gfp*) strains. Presented are overlay images of colorimetric channel (blue) with fluorescence from GFP (green), captured at the indicated time points post incubation using ChemiDoc MP imaging system. Shown are full-plate images, including both the antibiotic-free and antibiotic-containing regions (Cm+Spec), with white lines demarcating the border between the two.

**Table S1:**
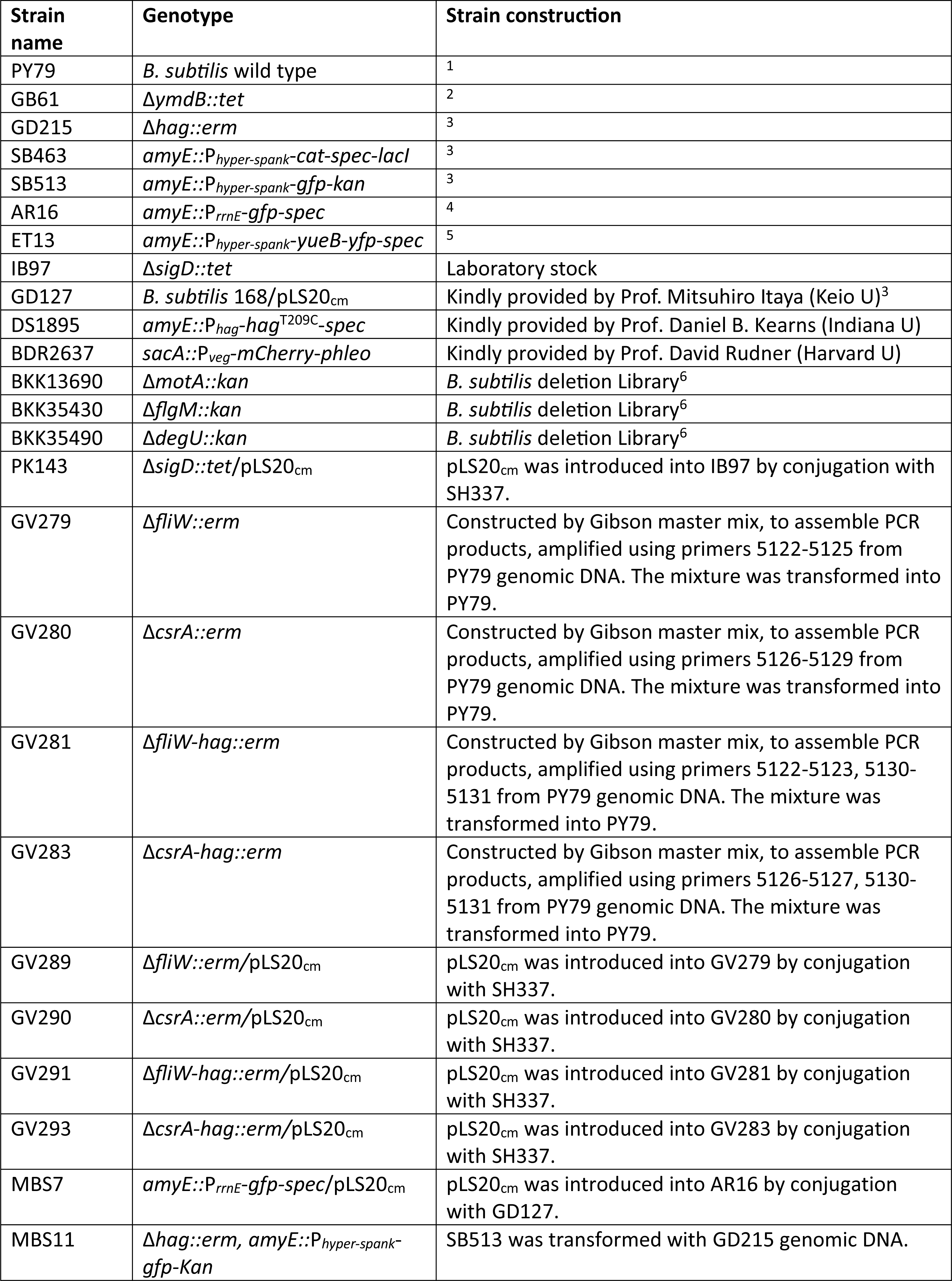
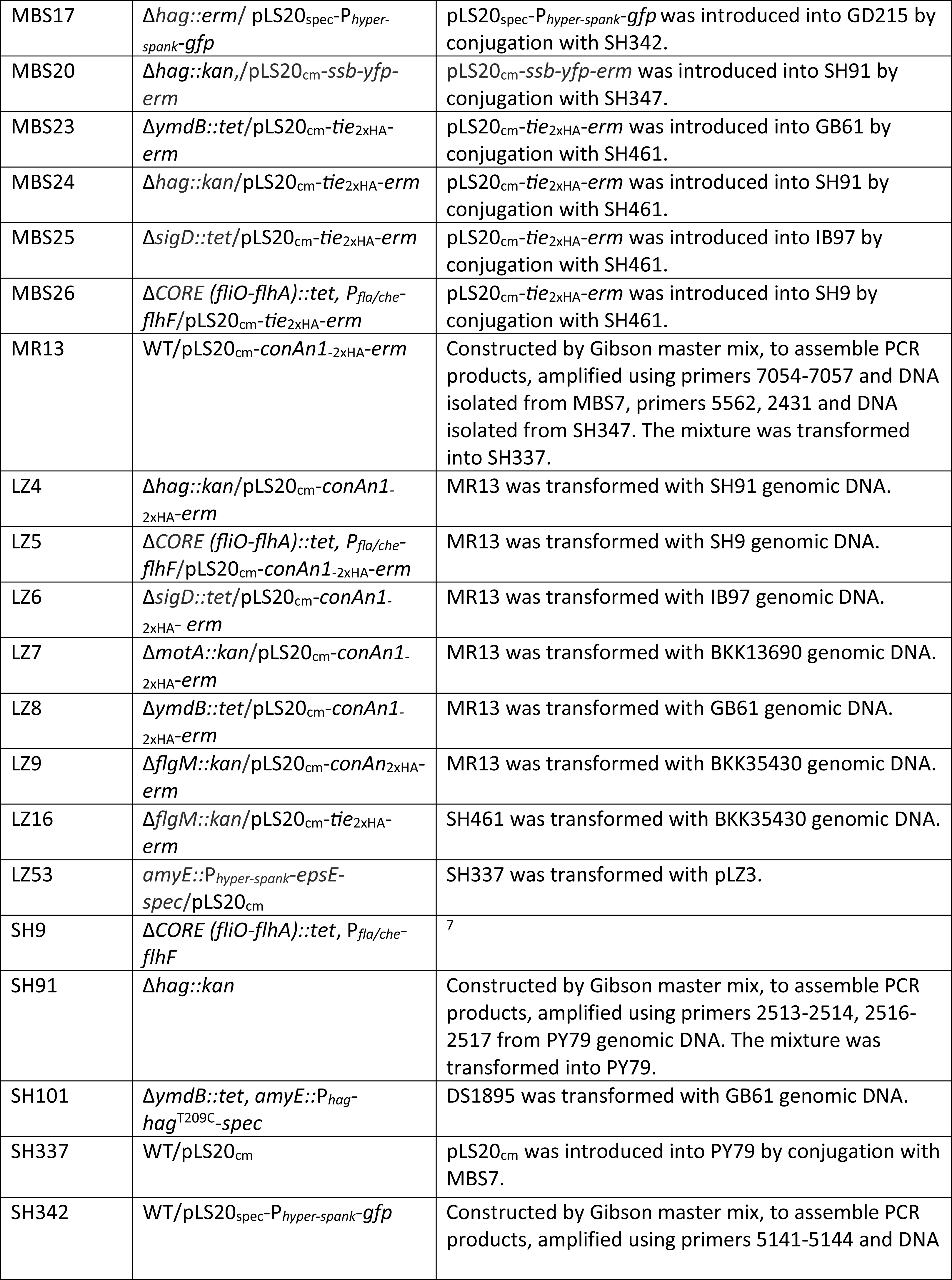
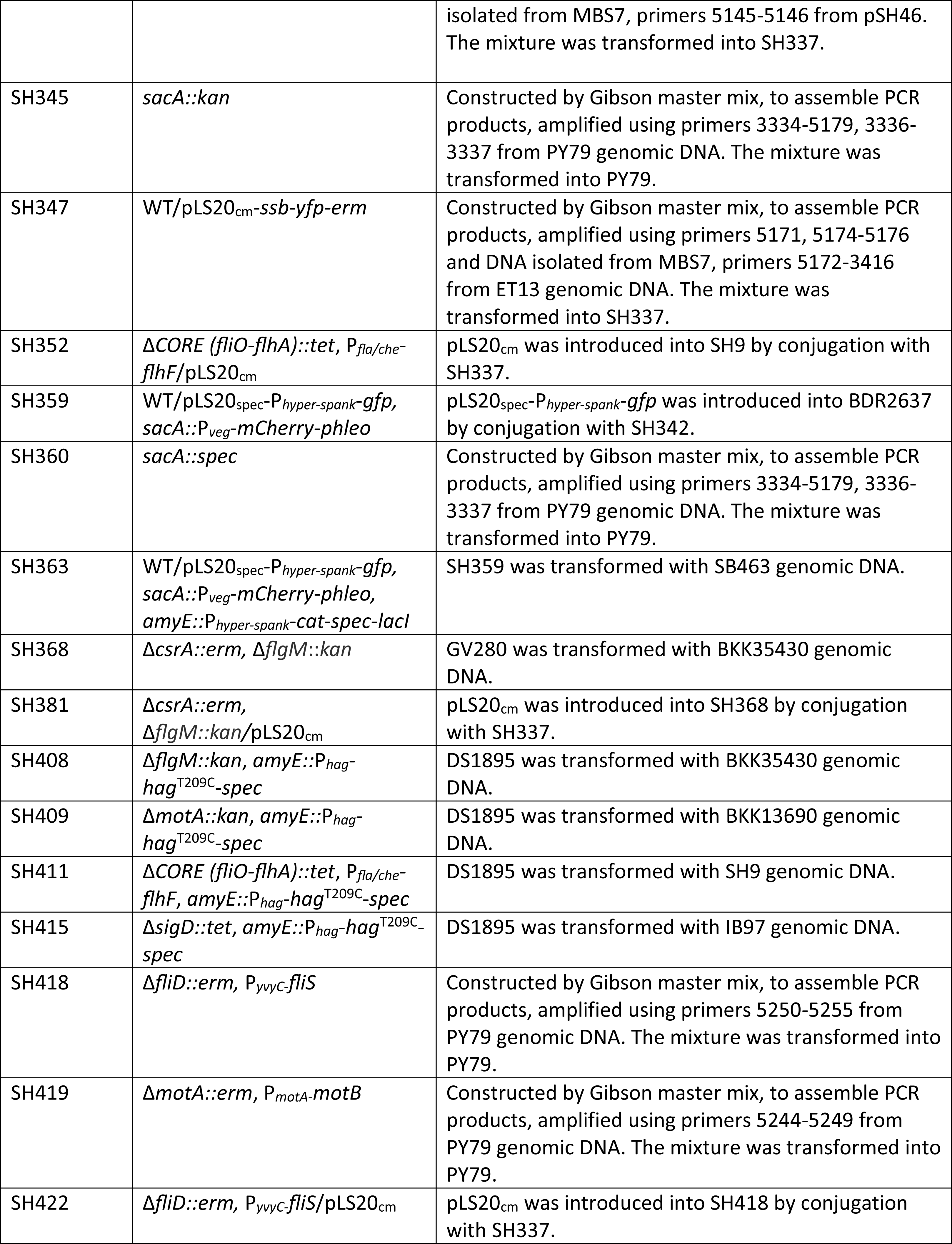
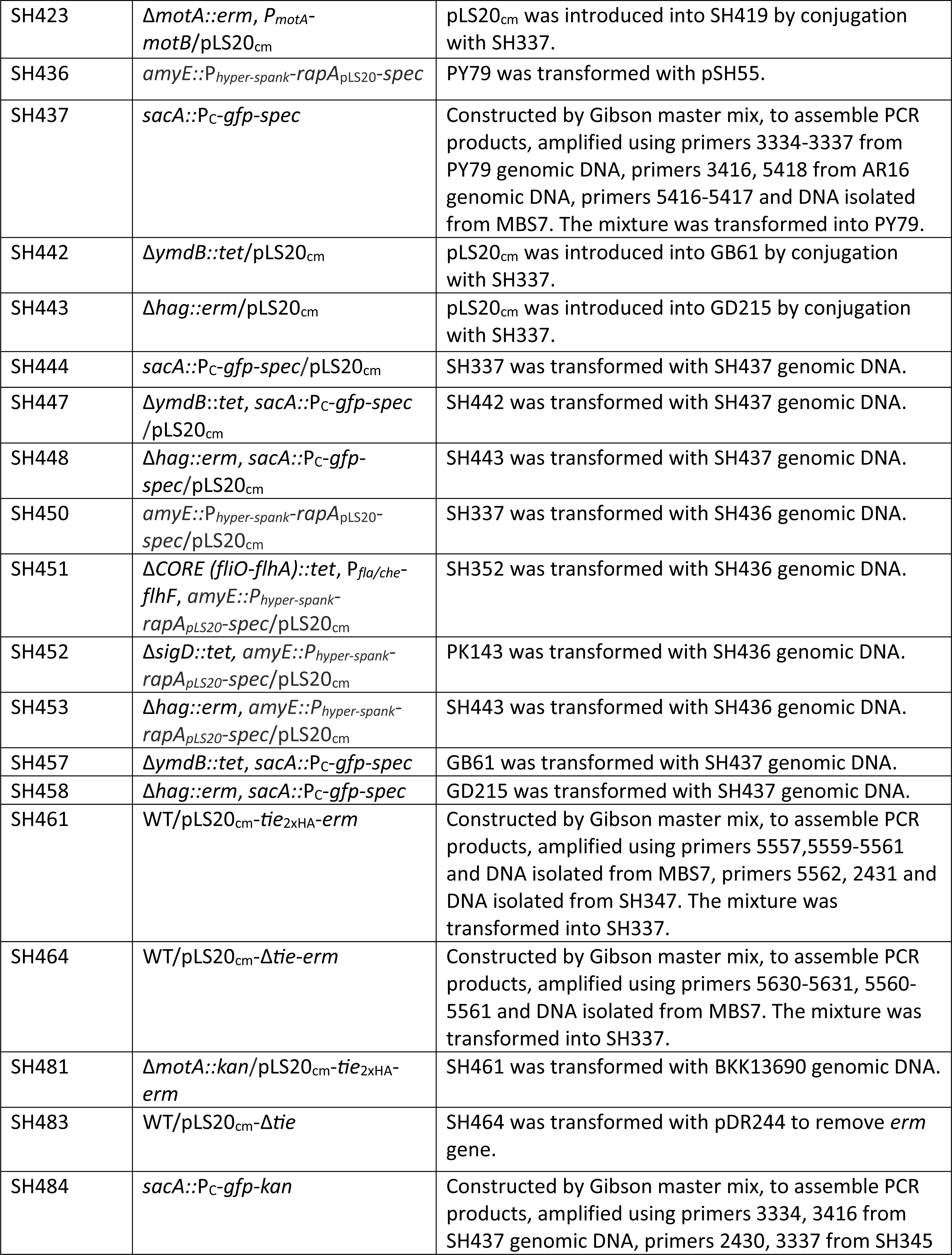
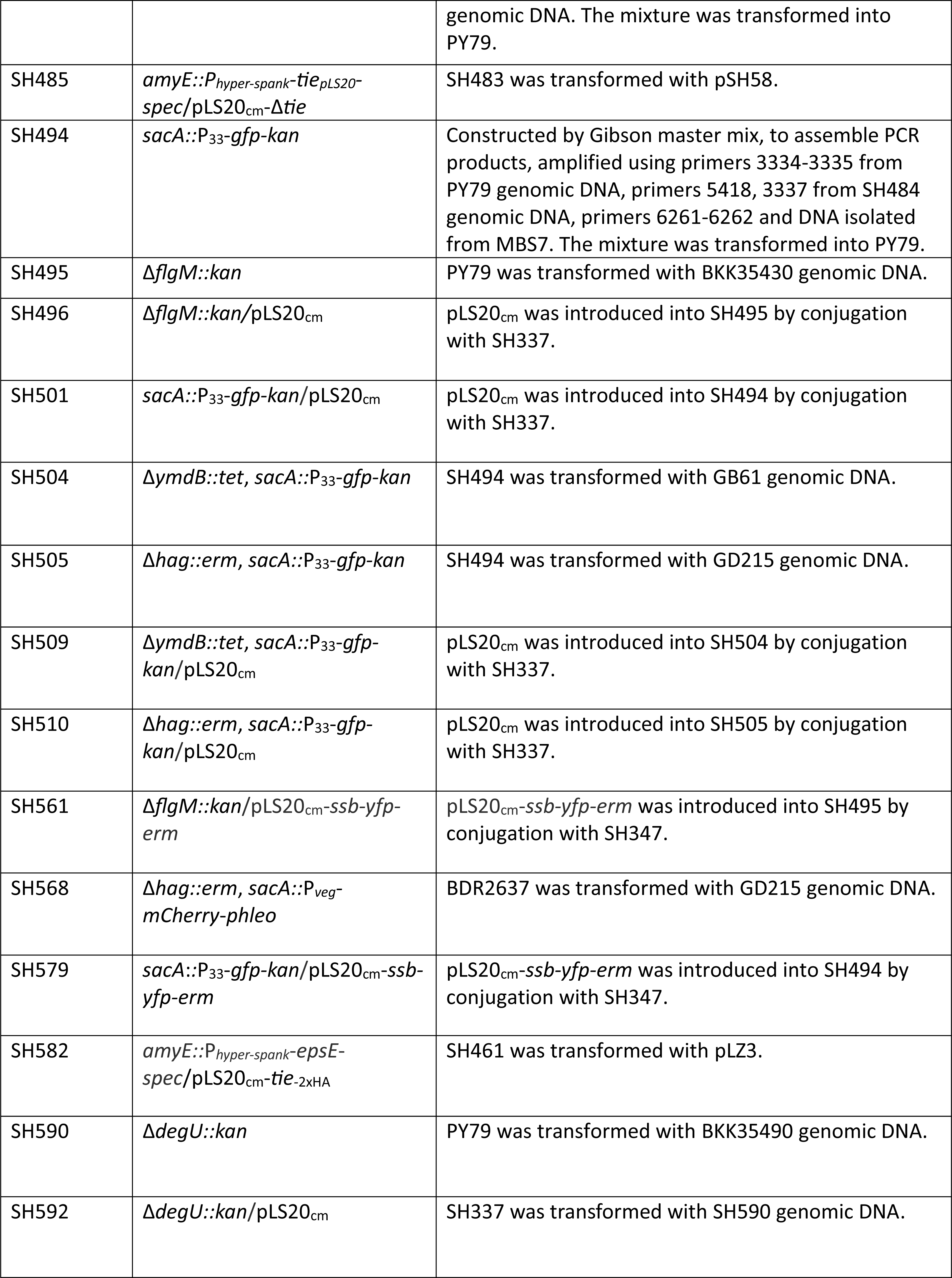
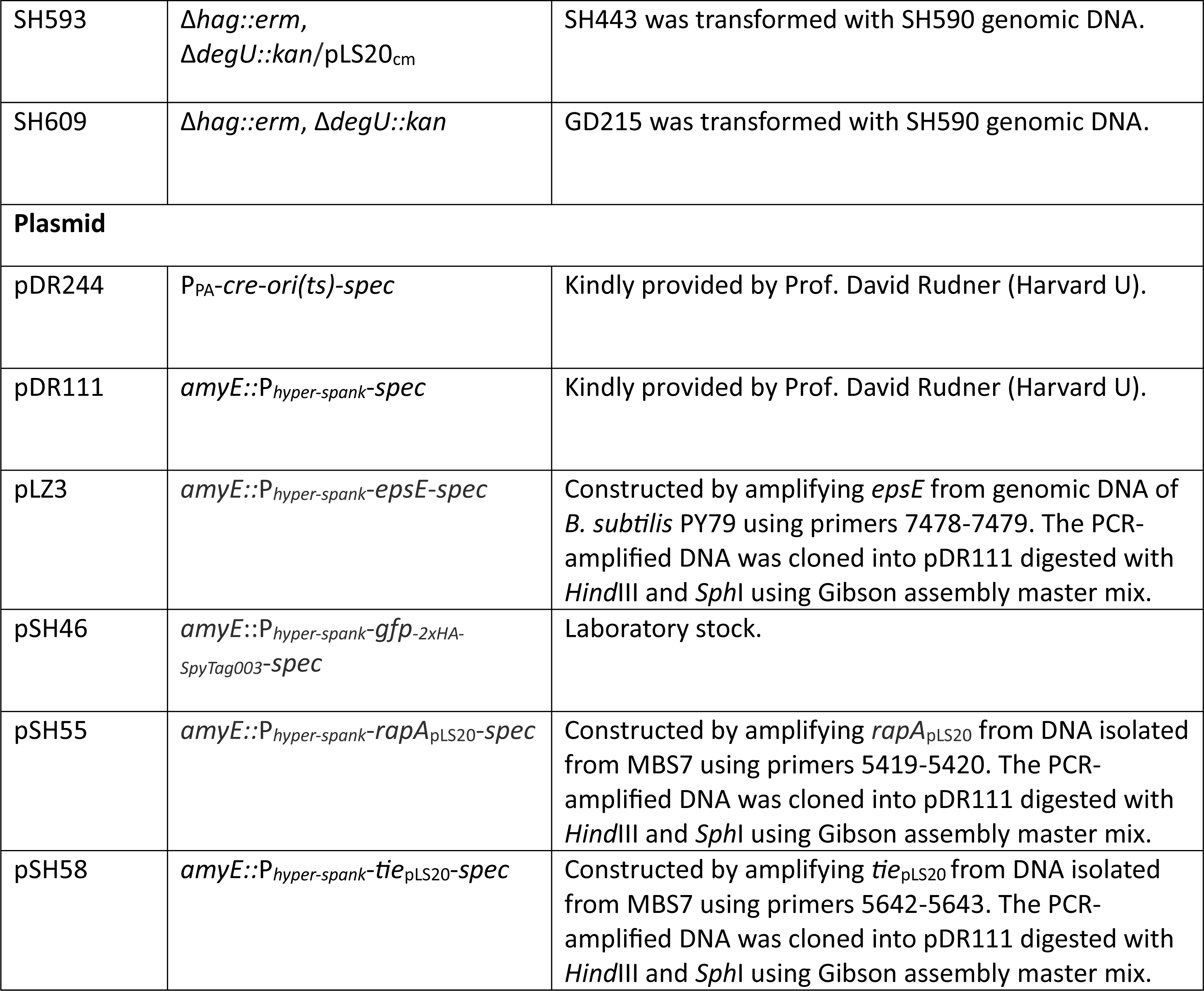
List of bacterial strains and plasmids used in this study.

**Table S2:**
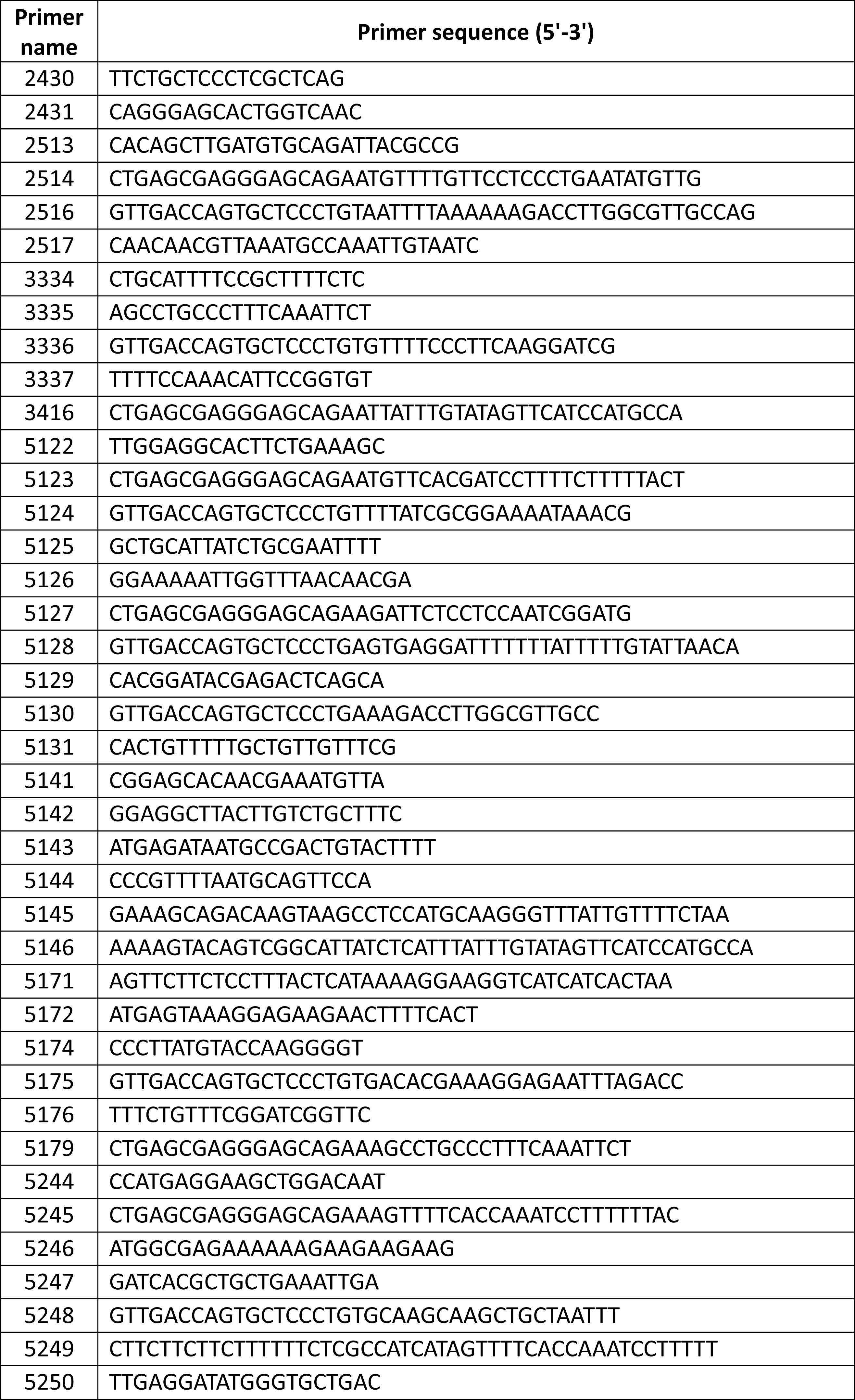
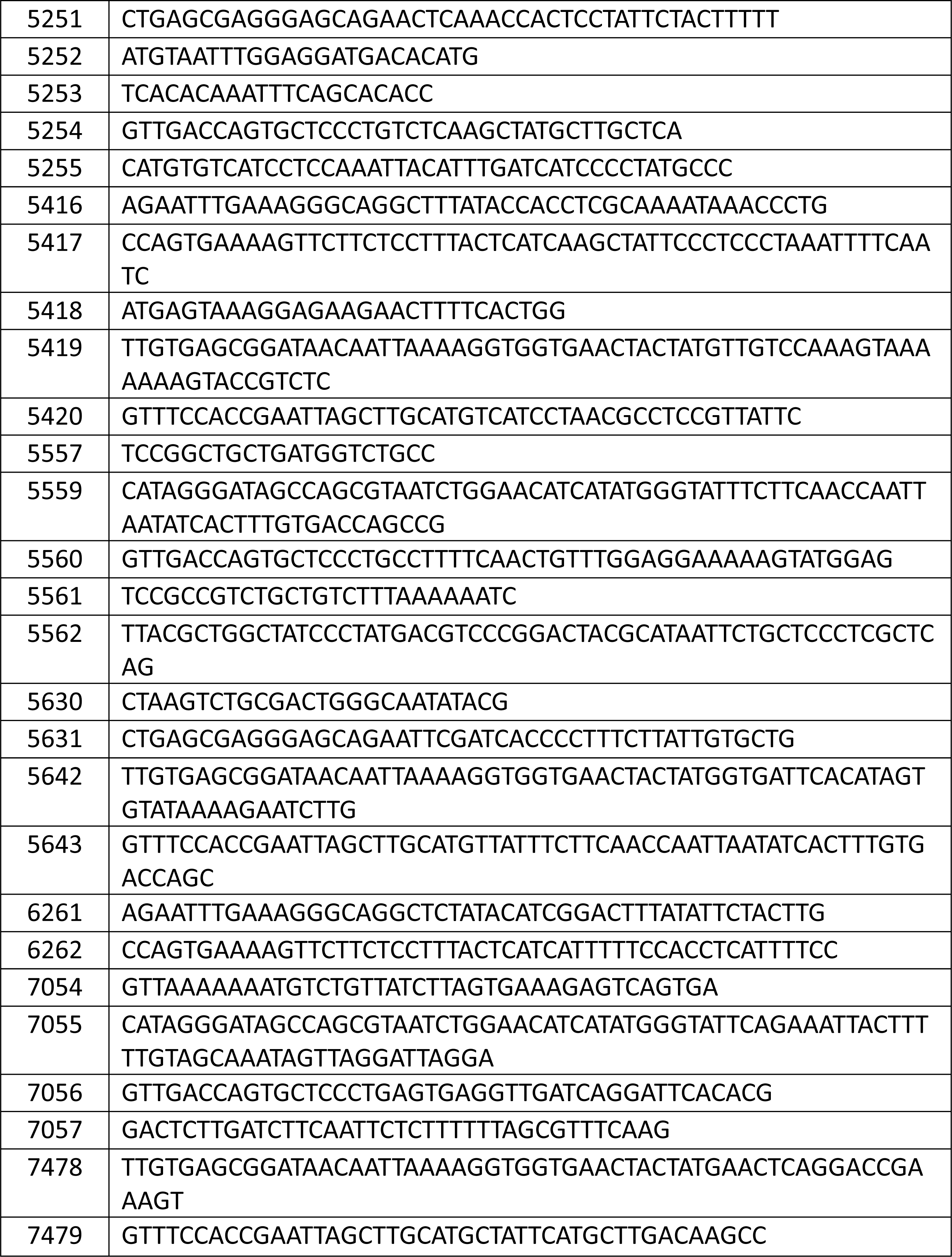
List of primers used in this study.

